# ESX-1 remodels the *Mycobacterium tuberculosis* antigen storage compartment in macrophages

**DOI:** 10.64898/2026.10.03.756392

**Authors:** Chak Hon Luk, Tania Auchynnikava, Antony Fearns, Julio Ortiz Canseco, Parise K. Lockwood, Camille Charoy, Maximiliano G. Gutierrez

## Abstract

*Mycobacterium tuberculosis* (Mtb) secretes antigenic proteins that drive protective T cell responses, yet how Mtb intracellular localization affects the trafficking and spatiotemporal distribution of mycobacterial antigens in macrophages remains elusive. Here, we developed a split fluorescent protein reporter to monitor the dynamics of the TB vaccine lead antigen Ag85B in infected human macrophages. We discovered that Mtb triggers the formation and remodeling of two distinct types of Mtb Antigen Storage Compartment (MASC) that retains Ag85 within macrophage membranes. Using subcellular spatial proteomics, we profiled the composition of the MASC, identifying both the Mtb and macrophage proteins that constitute these compartments. Unexpectedly, MASCs are defined by metabolic compartmentalization and rewiring to lipid metabolism in both the macrophage and Mtb. Thus, Mtb builds via the ESX-1 system a metabolically distinct antigen storage compartment in immune cells, with potential implications for antigen selection and delivery in TB vaccine development.

## Introduction

Tuberculosis (TB), caused by *Mycobacterium tuberculosis* (Mtb) is the leading cause of death by a single pathogen, leading to 10.7 million incident cases and 1.23 million deaths in 2024^1^. The Bacillus Calmette-Guérin (BCG) vaccine against TB was developed by Albert Calmette and Camille Guérin at Institut Pasteur and first administered to humans in 1921^2^. The BCG vaccine is effective in providing protection against severe forms of childhood TB, yet the protection fades as individuals reach adulthood^3^.

The efforts in improving BCG vaccines converge on two major strategies, firstly, the delivery of exogenous Mtb antigens with adjuvants and secondly, the expression of Mtb antigens via delivery of antigen-encoding nucleic acid or viral vectors. These efforts have yielded limited improvement to the existing protection of the current BCG vaccine^4^, warranting questions on what makes a TB vaccine effective, in addition to the mere presence of the antigens in the body. Mtb has several identified antigens that are being used as biomarkers for TB diagnosis as well as candidates for vaccine development. These Mtb antigens can be classified by their biochemical nature or cellular localization and function, yet there is a limited understanding of how these Mtb antigens elicit protective immunity in humans.

Mtb antigens encompass protein, lipoprotein, lipid and carbohydrate antigens. While several protein antigens are widely used in TB diagnosis^5^, their subcellular localization in human cells remains poorly characterized^6,7^ in the context of the mycobacterial envelope^8,9^, lipid^10–12^, glycolipid^10,13^ and lipoprotein^14^ antigens. This knowledge gap is primarily due to the lack of tools for Mtb protein antigen visualization and the limited efficient usage of bulky tags that will affect protein secretion/export. Antibodies have been successfully used to detect some Mtb secreted proteins, with only a handful fully validated using knockout mutants for imaging^15,16^. It is also clear that only a subpopulation of intracellular Mtb secretes the antigens as observed for Ag85 in postmortem tissues^17^. This heterogenous secretion of Mtb antigens cannot be captured using bulk assays^18,19^, and have precluded our understanding of how much secretion of Mtb antigens occur at the single-cell level during early infection.

The subcellular localization of bacterial antigens plays a key role in determining how they are processed and presented to distinct subsets of T cells. Antigens in the phagolysosome are degraded by lysosomal proteases and loaded onto Major Histocompatibility Complex (MHC) Class II for presentation to CD4+ T cells. Simultaneously, antigens that are expressed or released into the cytosol are instead degraded by the proteasome, transported to the endoplasmic reticulum by TAP and loaded onto MHC Class I for recognition by CD8+ T cells^20^. Mtb resides within membrane-bound phagolysosomal compartments but also can damage the phagosomal membrane and access the cytosol through the Type VII secretion system ESX-1 and lipids^21^. In addition to cytosolic access, Mtb triggers via the ESX-1system the formation of complex endocytic membrane structures in its vicinity ^22–24^. Although ESX-1 activity has been shown to be required for MHC-I presentation of peptides derived from T7SS substrates^25^, much remains unknown about where Mtb antigens are localized and how they are processed, given the heterogeneity of Mtb intracellular localization.

The Ag85 complex, comprised of three isoforms (A, B and C in stoichiometry of 2:3:1) is the most abundant secreted protein antigen among mycobacteria species. Ag85 catalyzes the biosynthesis of essential components of the bacterial envelope^26–28^. In infected cells, the Ag85 complex is secreted by the bacterium within the phagosome and localizes to both the Mtb outer membrane and host endocytic compartments^29^. However, the fate of secreted Ag85 from cytosolic Mtb remains unknown.

Here, we report the generation of a split fluorescent reporter system to visualize the spatiotemporal dynamics of Ag85B secretion during Mtb infection of human macrophages. By monitoring the subcellular localization of tagged and endogenous Ag85, we found that Ag85 reaches the cytosol of infected macrophages via the ESX-1 secretion system. Ag85 not only accessed the cytosol but was predominantly retained in a compartment with distinct unappreciated membranous structures which we term the Mtb Antigen Storage Compartment (MASC). Using spatial proteomics, we discovered that the MASC undergoes striking ESX-1-dependent metabolic remodeling, uncovering ESX-1 as a novel player in the Mtb metabolic adaptation to the host cell. Together, these data provide evidence for the existence of a Mtb antigen storage compartment that is regulated by ESX-1 activity.

## Results

### Ag85B is translocated from the Mtb phagosome to the macrophage cytosol

To define the spatiotemporal localization of Ag85B secreted by intracellular Mtb in human macrophages, we devised a two-component fluorescent reporter system^30^ (**Figure 1A**) based on a strong split mNeonGreen (mNG) fluorescence protein^31^ and signal amplification with tandem repeats. This system is comprised of the first 10 strands of mNeonGreen3K (mNG_1-10_) and seven repeats of the complementary 11^th^ strand of mNeonGreen3K (7xmNG_11_). We generated a Mtb H37Rv strain constitutively expressing E2-Crimson and Ag85B-7xmNG_11_, fused to the 7xmNG_11_. On the other hand, we generated THP-1 macrophages constitutively expressing mNG_1-10_ in the cytosol (**Figure 1A**). We then infected THP-1 mNG_1-10_ macrophages with Mtb H37Rv Ag85B-7xmNG_11_. After 2 h of infection, only background fluorescence was observed in infected cells (**Figure 1B** and **Supp. Figure 1A and 1B**). In contrast, macrophages infected with Mtb H37Rv Ag85B-7xmNG_11_ showed mNG signal in intracellular bacteria and the cytosol after 72 h of infection (**Figure 1B and Supp. Figure 1A**). This signal was specific as THP-1 mNG_1-10_ infected with Mtb expressing only E2-Crimson or THP-1 macrophages infected with Mtb H37Rv Ag85B-7xmNG_11_ only showed background fluorescence (**Supp. Figure 1A and 1B).** Quantitatively, there was an increase in both the percentage of mNG+ intracellular Mtb at 72 h (**Figure 1C)** and the percentage of mNG+ macrophages compared to uninfected cells (**Figure 1D**). Confirming the imaging analysis, a flow cytometry analysis showed an increase in the number of mNG+ macrophages from 2 to 72 h after infection (**Figure 1E** and **Supp. Figure 2A and 2B**). Altogether, we established a reporter to monitor antigen trafficking and found that Ag85B is secreted from the phagosomes into the cytosol of infected macrophages.

**Figure 1.**
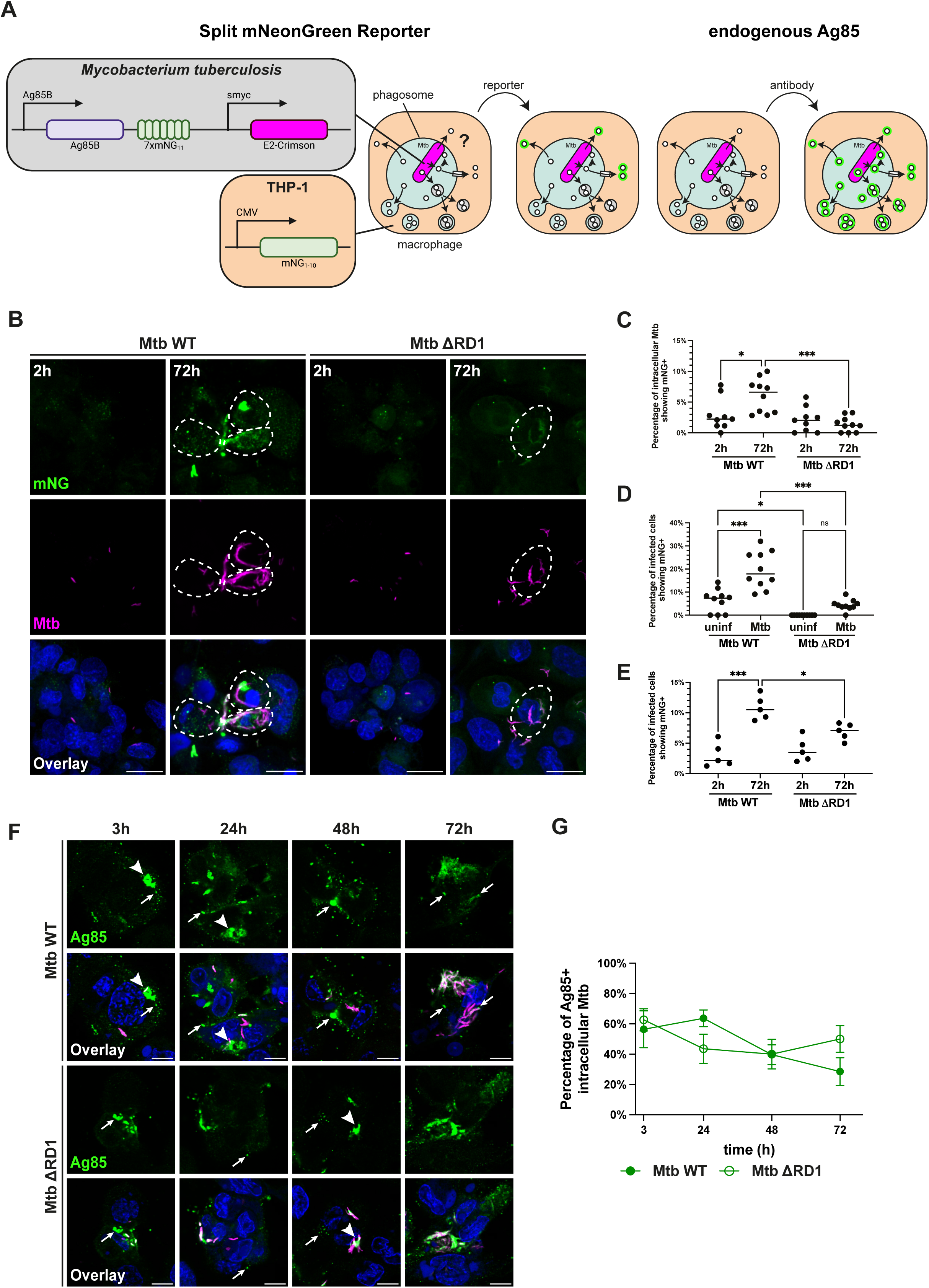
Ag85B accumulates in an antigen storage compartment and translocates to the cytosol via the ESX-1 secretion system. (A) Schematic of labelling mechanism of Mtb Ag85B split fluorescent reporter and antibody staining to monitor antigen localization. (B) Representative confocal images of Mtb H37Rv WT and ΔRD1 Ag85B-7xmNG_11_ infected THP-1 mNG_1-10_ at 3h and 72h pi, images showing nuclei (blue), mNG (green) and Mtb (magenta). Scalebar = 20 µm. (C) Quantification of mNG+ Mtb in THP-1 mNG_1-10_ cells infected by Mtb H37Rv WT and ΔRD1 Ag85B-7xmNG_11_ at 3h and 72h pi by microscopy. (D) Quantification of mNG+ THP-1 mNG_1-10_ cells infected by Mtb H37Rv WT and ΔRD1 Ag85B-7xmNG_11_ at 3h and 72h pi by microscopy. (E) Quantification of mNG+ THP-1 mNG_1-10_ cells infected by Mtb H37Rv WT and ΔRD1 Ag85B-7xmNG_11_ at 3h and 72h pi by flow cytometry. (F) Representative confocal images of THP-1 cells infected by Mtb H37Rv WT and ΔRD1 at 3h, 24h, 48h and 72h pi, images showing nuclei (blue), Ag85 (green) and Mtb (magenta), small vesicles are depicted by arrows and TVS are depicted by arrowheads. Scalebar = 10 µm. (G) Quantification of Ag85+ Mtb in THP-1 cells infected by Mtb H37Rv WT and ΔRD1 at 3h, 24h, 48h and 72h pi.

**Figure 2.**
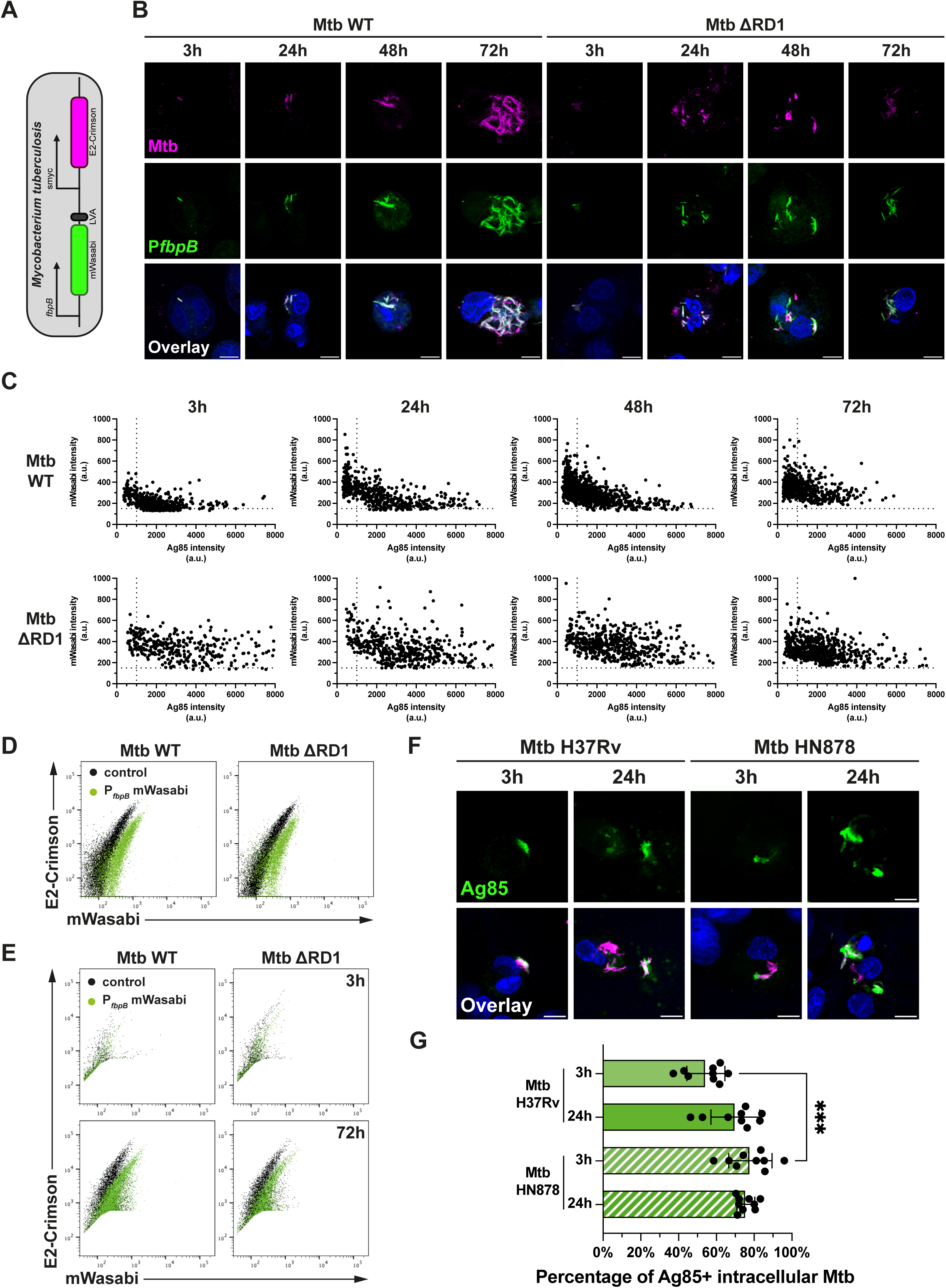
Ag85 localization in macrophages is not associated with expression levels. (A) Schematic of Ag85B transcription reporter. (B) Representative confocal images of THP-1 cells infected by Mtb H37Rv WT and ΔRD1 with P*fbpB*-mWasabi-LVA reporter at 3h, 24h, 48h and 72h pi, images showing nuclei (blue), P*fbpB*-mWasabi-LVA (green) and Mtb (magenta). (C) Scatter plot of P*fpbB*-mWasabi-LVA intensity against Ag85 intensity of Mtb H37Rv WT and ΔRD1 in THP-1 cells at 3h, 24h, 48h and 72h pi. (D) Flow cytometry analysis of Mtb H37Rv WT and ΔRD1 inoculum; control (black) and P*fbpB*-mWasabi-LVA reporter (green). (E) Flow cytometry analysis of THP-1 cells infected by Mtb H37Rv WT and ΔRD1 P*fbpB*-mWasabi-LVA reporter at 3h and 72h pi; control (black) and P*fbpB*-mWasabi-LVA reporter (green). (F) Representative confocal images of THP-1 cells infected by Mtb H37Rv WT and Mtb HN878 WT at 3h and 24h pi, images showing nuclei (blue), Ag85 (green) and Mtb (magenta). Scalebar = 10 µm. (G) Quantification of Ag85+ Mtb in THP-1 cells infected by Mtb H37Rv WT and Mtb HN878 WT at 3h and 24h pi

### Ag85B translocation to the macrophage cytosol is ESX-1-dependent

Next, we focused on understanding how Ag85B reached the macrophage cytosol. Mtb triggers phagosome damage^21^ and we hypothesized that phagosome damage is required for Ag85 translocation to the cytosol. To test this, we generated Mtb H37Rv ΔRD1 Ag85B-7xmNG_11_, a mutant that does not induce membrane damage^21^. As Ag85B-7xmNG_11_ only gives a fluorescent signal after accessing the host cytosol and complementing the cytosolic mNG_1-10_, we used the split system to validate whether Ag85B accessed the host cytosol via phagosome membrane damage (**Figure 1A**). There was no significant increase in mNG+ signal in macrophages infected with Mtb ΔRD1 Ag85B-7xmNG_11_ when compared with Mtb WT Ag85B-7xmNG_11_ infected macrophages (**Figure 1B to 1D**). Analysis by flow cytometry confirmed that Mtb WT Ag85B-7xmNG_11_ infected macrophages showed a higher mNG+ signal than Mtb ΔRD1 Ag85B-7xmNG_11_ infected macrophages (**Figure 1E**), indicating the translocation of Ag85B to the host cytosol required a functional ESX-1 secretion system.

To validate these findings, we analyzed the localization of endogenous Ag85 (**Figure 1A**). We found that the commonly used antibody to detect Ag85B (RRID:AB_776575) also recognized the other isoforms Ag85A and Ag85C (**Supp. Fig 3A and 3B**). Therefore, when analyzing endogenous levels, we refer to all three isoforms as Ag85. We observed that after infection with Mtb WT, Ag85 associated with intracellular bacteria, localized in phagosomes, predominantly in vesicular compartments and throughout the cytosol. On the other hand, after infection with Mtb ΔRD1, endogenous Ag85 localized mostly with bacteria, phagosomes and vesicular compartments (**Figure 1F, arrows**). Previous studies have reported the presence of Ag85 in a class of export vesicles named antigen export vesicles (AEVs) present in antigen presenting cells^29^. To determine if these Ag85+ vesicles share similar identities to AEVs, we examined the association of Ag85+ vesicles with α-tubulin. We did not observe localization of α-tubulin to Ag85+ compartments, indicating these compartments were distinct from previously reported AEVs (**Supp. Figure 4A and B**). A quantitative analysis showed that the percentage of Ag85+ bacteria was similar after 3 h of infection in both Mtb WT and Mtb ΔRD1 but decreased with time in Mtb WT infected macrophages (**Figure 1G).** Altogether, these data show that Ag85 reaches the cytosol in an ESX-1-dependent manner, while both Mtb WT and Mtb ΔRD1 secrete Ag85 into the phagosome and vesicular compartments that we named Mtb antigen storage compartment (MASC) hereafter.

**Figure 3.**
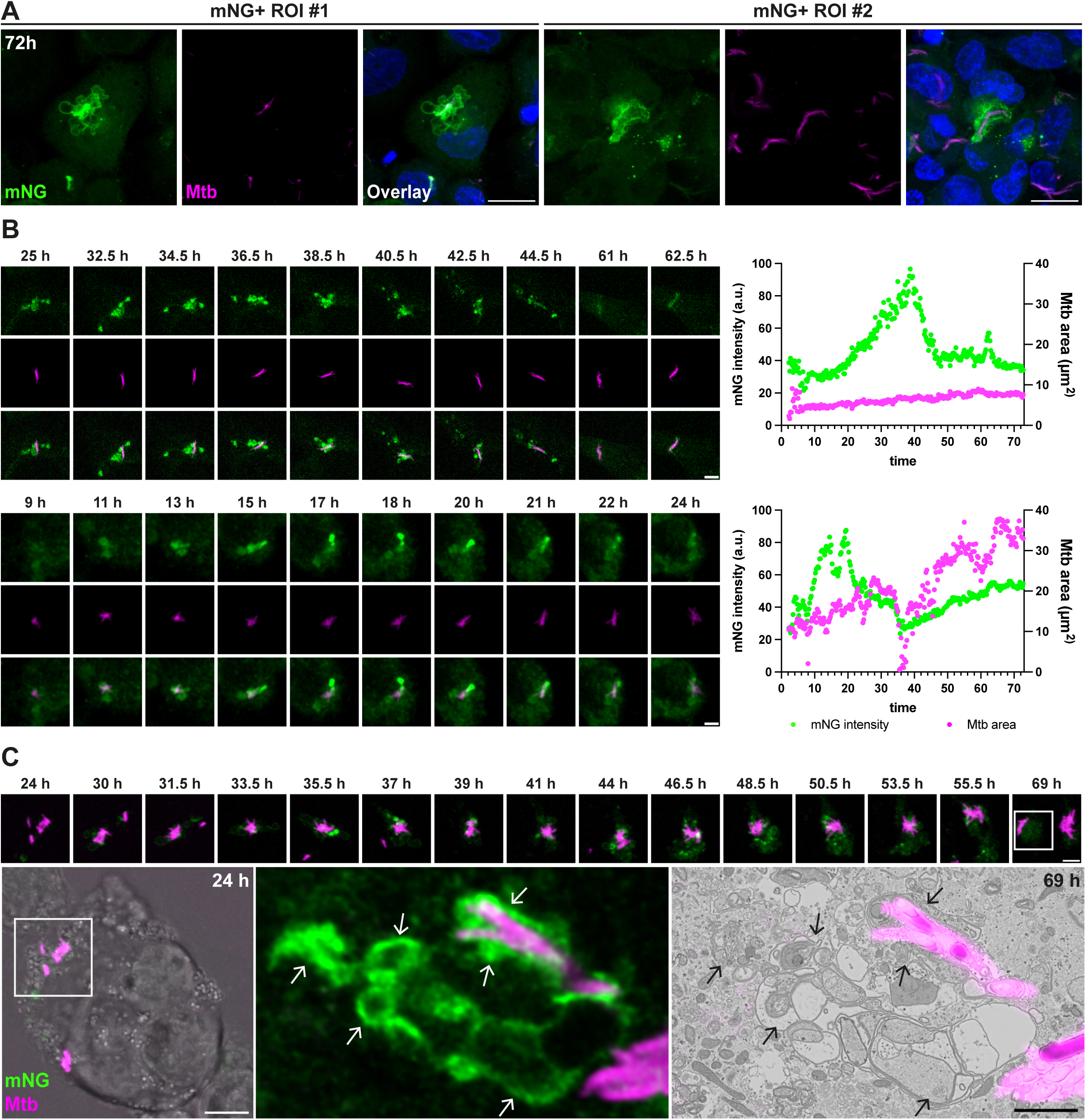
Complemented Ag85B split reporter associates with the membranes of the antigen storage compartment in macrophages. (A) Representative confocal images of Mtb H37Rv WT Ag85B-7xmNG_11_ infected THP-1 mNG_1-10_ at 72h pi, images showing nuclei (blue), mNG (green) and Mtb (magenta). Scalebar = 20 µm. (B) Live microscopy snapshots and fluorescence intensity quantification of intracellular Mtb shed or encapsulated by complex membrane structures, images showing secreted Ag85B-7xmNG_11_ (green) and Mtb (magenta). Scalebar = 5 µm. (C) Live-CLEM snapshots of intracellular Mtb, images showing mNG (green) and Mtb (magenta). Scalebar = 5 µm. Transmission electron micrograph showing the Ag85+ structures (indicated by arrows) proximal to Mtb at 69h pi. Scalebar = 2 µm.

**Figure 4.**
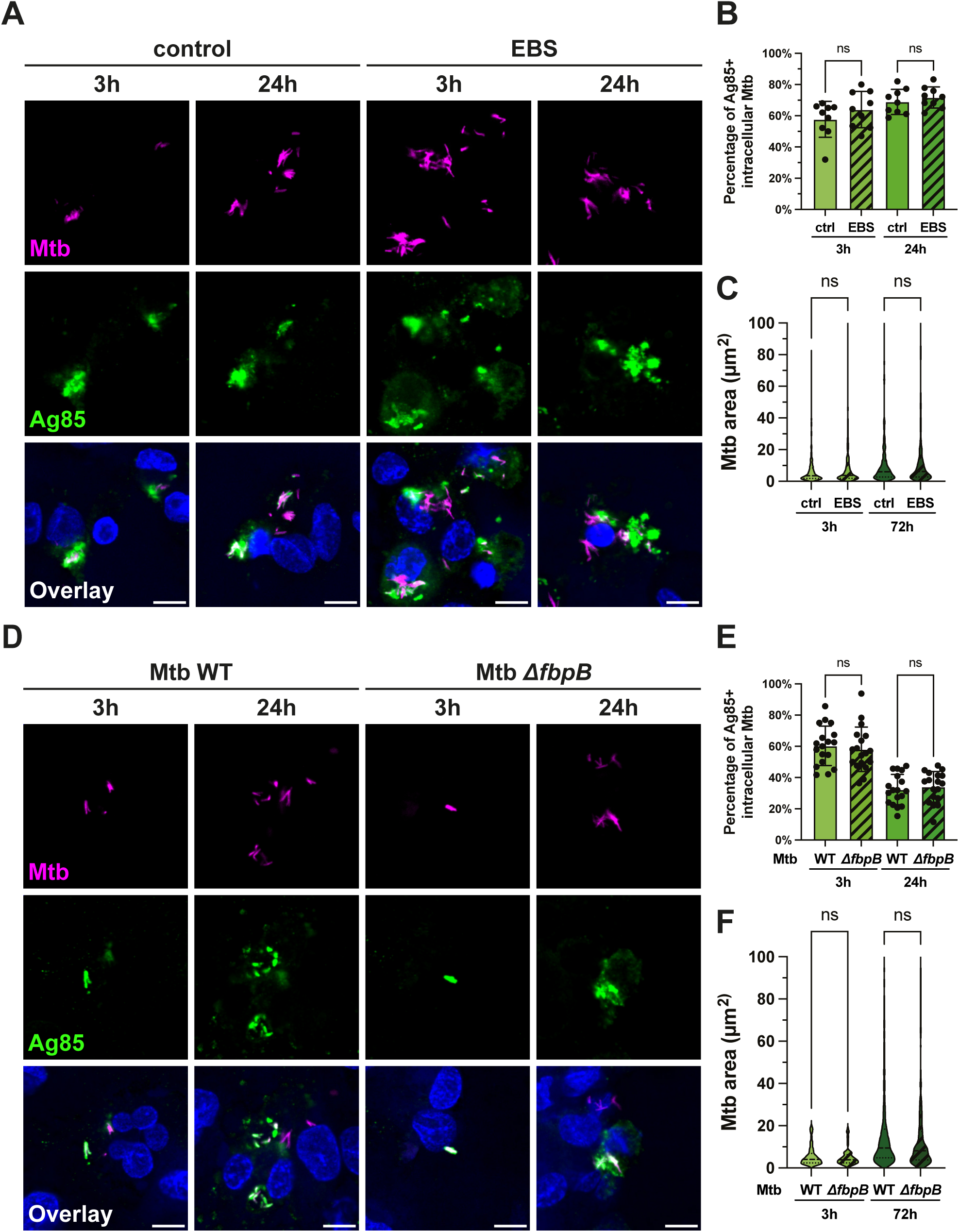
Ag85 activity is dispensable for the Ag85 storage compartment in macrophages. (A) Representative confocal images of THP-1 cells infected by control and EBS-treated Mtb H37Rv WT at 3h and 24h pi, images showing nuclei (blue), Ag85 (green) and Mtb (magenta). Scalebar = 10 µm. (B) Quantification of Ag85+ Mtb in THP-1 cells infected by control and EBS-treated Mtb H37Rv WT at 3h and 24h pi. (C) Quantification of Mtb area in THP-1 cells infected by control and EBS-treated Mtb H37Rv WT at 3h and 72h pi. (D) Representative confocal images of THP-1 cells infected by Mtb H37Rv WT and *ΔfbpB* at 3h and 24h pi, images showing nuclei (blue), Ag85 (green) and Mtb (magenta). Scalebar = 10 µm. (E) Quantification of Ag85+ Mtb in THP-1 cells infected by Mtb H37Rv WT and *ΔfbpB* at 3h and 24h pi. (F) Quantification of Mtb area in THP-1 cells infected by Mtb H37Rv WT and *ΔfbpB* at 3h and 72h pi.

### Both Mtb WT and Mtb ΔRD1 express similar levels of Ag85 in infected macrophages

We observed MASC in a fraction of macrophages infected by Mtb WT and Mtb RD1. One possible interpretation is that the Ag85 expression after infection was different at the single macrophage level. To test this, we used a dual transcriptional reporter using mWasabi fused to a degradation tag^32^ (Aylan et al., *in press*) to analyze the promoter activity of *fbpB* as a proxy of Ag85B production (**Figure 2A**). Using this transcriptional reporter, we observed a homogenous *fbpB* promoter activity in Mtb before and after infection by both imaging (**Figure 2B and 2C**) and flow cytometry (**Figure 2D and 2E**). The Mtb HN878 strain shows higher levels of Ag85 secretion than Mtb H37Rv despite comparable gene expression^18,19^. When we measured the incidence of Ag85+ MASC in Mtb H37Rv and Mtb HN878 infected macrophages, we observed Ag85+ MASC in both Mtb strains. However, Mtb HN878 showed an increased number of Ag85+ MASC at 3 h after infection as compared to H37Rv (**Figure 2F and 2G**). Altogether, these results suggest that the formation of Ag85+ MASC is not associated with differential levels of expression.

### Ag85 translocation via cytosol of infected cells to uninfected bystander cells is ESX-1 dependent

Using the split fluorescent reporter and antibody staining to depict the localization and historical path of Ag85, we further applied them to understand the Ag85 localization in uninfected macrophages (**Figure 1A** and **Supp. Figure 5A**). Using the split reporter, we observed that the mNG signal was not only present in infected cells but also in uninfected cells (**Supp. Figure 5B**). A quantitative analysis showed that the number of uninfected THP-1 mNG_1-10_ cells positive for mNG increased during the infection in an ESX-1-dependent manner (**Supp. Figure 5C**). This observation suggests that cytosolic Ag85B can be transferred to uninfected cells and confirms the function of ESX-1 in Ag85 cytosolic localization (**Supp. Figure 5C**). To validate these findings, we analyzed the presence of endogenous Ag85 among uninfected cells from an infected sample (bystander). We detected Ag85 localized in host endocytic vesicles in bystander cells (**Supp. Figure 5D**). Moreover, we observed an elevated fraction of Ag85+ bystander cells at 48 h after infection, suggesting that cell-to-cell transfer takes place as the infection progressed (**Supp. Figure 5E**). In contrast to the readout from the split fluorescent reporter signal, both infection with Mtb WT and Mtb ΔRD1 resulted in Ag85+ bystander macrophages (**Supp. Figure 5D and E**). Altogether, these data show that the access of Ag85 to the cytosol is not a requirement for cell-to-cell transfer, where Ag85 can be transferred to bystander macrophages through different paths.

**Figure 5.**
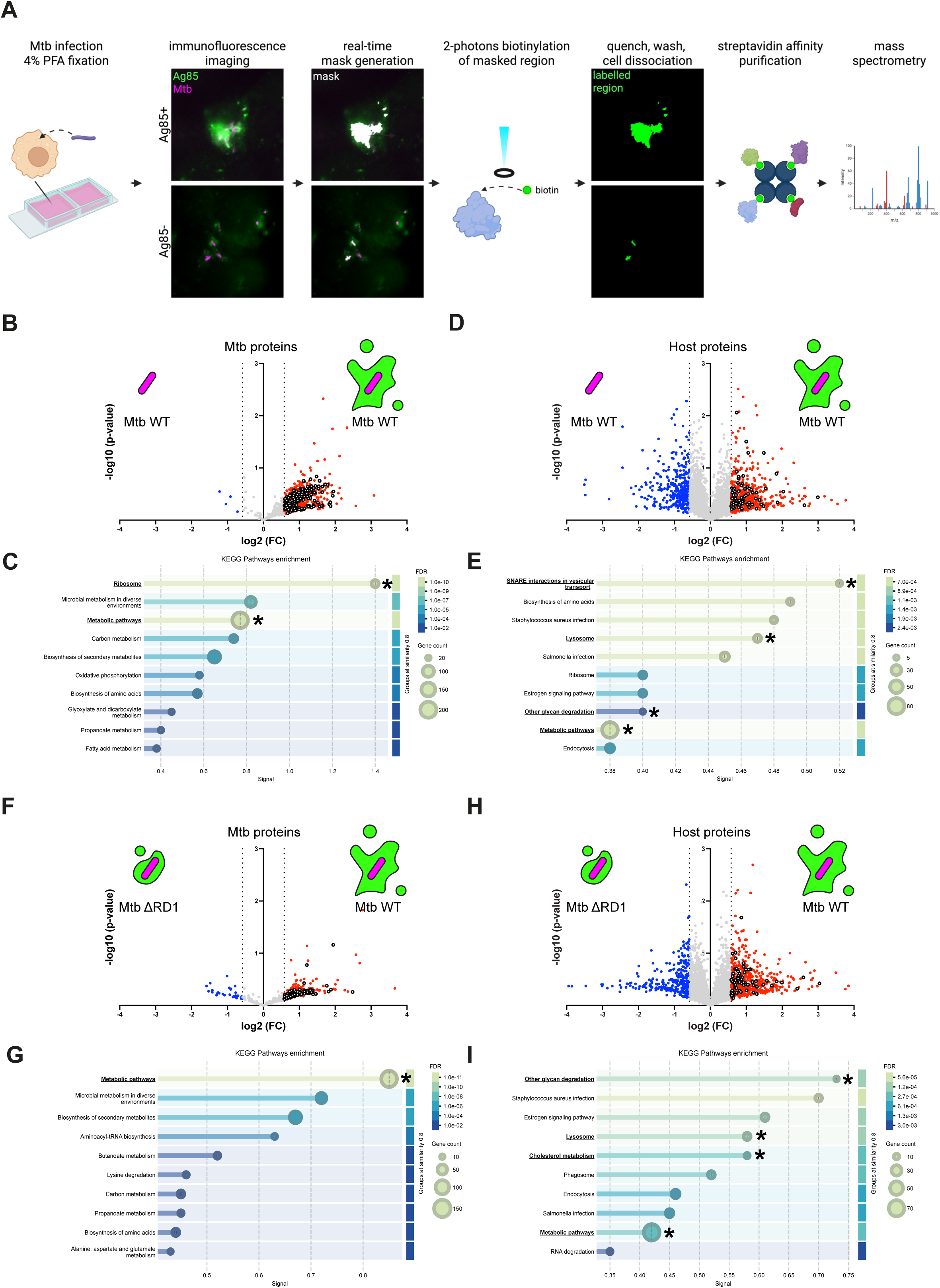
The Ag85 storage compartment is a metabolic hub rewired by the ESX-1 secretion system. (A) Schematic diagram of spatial proteomic workflow. (B) Pairwise comparison of enriched and depleted Mtb proteins in Ag85+ structures relative to Ag85- Mtb induced by Mtb-WT at 3h after infection. Enriched proteins of metabolic pathways from KEGG pathway analysis are highlighted. (C) KEGG pathway analysis of enriched proteins in (B). (D) Pairwise comparison of enriched and depleted host proteins in Ag85+ structures relative to Ag85- Mtb induced by Mtb-WT at 3h after infection. Enriched proteins of metabolic pathways from KEGG pathway analysis are highlighted. (E) KEGG pathway analysis of enriched proteins in (D). (F) Pairwise comparison of enriched and depleted Mtb proteins in Ag85+ structures induced by Mtb-WT relative to Mtb-ΔRD1 at 3h after infection. Enriched proteins of metabolic pathways from KEGG pathway analysis are highlighted. (G) KEGG pathway analysis of enriched proteins in (F). (H) Pairwise comparison of enriched and depleted host proteins in Ag85+ structures induced by Mtb-WT relative to Mtb-ΔRD1 at 3h after infection. Enriched proteins of metabolic pathways from KEGG pathway analysis are highlighted. (I) KEGG pathway analysis of enriched proteins in (H). Plots showing –log_10_(p value) from two-sided Student’s t test and log_2_FC differences between each comparison.

### Ag85B associates with host membranes of the antigen storage compartment

Ag85 complex is highly abundant and localizes onto the outer membrane of replicating Mtb^26,27^. We detected endogenous Ag85 in Mtb phagosomes and the lumen of endocytic compartments (**Figure 1F**) as previously reported^7,29^. Using the split fluorescent reporter system, we found that the complemented mNG signal was not only in the lumen of the newly formed MASC, but also potentially associated with host membranes (**Figure 3A**). Using live microscopy to analyze the spatiotemporal dynamics of Ag85B-7xmNG_11_ localization in macrophages, we observed that through time Ag85B-7xmNG_11_ localized to both the membranes and lumen of the antigen storage compartment (**Figure 3B, Movie S1 and 2**). To confirm the membrane localization of Ag85B, we performed live-correlative light electron microscopy (CLEM) to localize the complementing Ag85B-7xmNG_11_ localization in infected macrophages with spatiotemporal resolution. By analyzing the ultrastructural details of the mNG+ membrane structures by CLEM, we confirmed that Ag85B-7xmNG_11_ localized in complex multi-membrane structures showing resemblance to the tubulovesicular structures previously reported^22–24^ and distinct from previously reported Ag85+ endocytic compartments^29^ (**Figure 3C and Movie S3**). Altogether, these data show that MASC can encompass intact and damaged phagolysosomal compartments and that Ag85 associates with host cell membranes of the damaged MASC.

### Ag85 activity is dispensable for the generation of the antigen storage compartment

We next asked whether Ag85 functionally contributes to MASC formation or is passively retained as cargo. Ag85 acts to remodel outer membrane surface and has a role in synthesizing Trehalose-6,6’-dimycolate (TDM)^33^ on the mycobacterial surface. Loss of Ag85A leads to disruption of plasma membrane domains and mycolic acid composition^34^. To understand if the Ag85+ MASC were caused by Ag85-mediated outer membrane remodeling of Mtb, we blocked Ag85 activity using ebselen (EBS) and measured the prevalence of Ag85+ MASC^35^. We observed similar numbers of Ag85+ MASC in control and EBS-treated conditions (**Figure 4A and 4B**). We also observed comparable levels of bacterial replication between EBS-treated and control conditions (**Figure 4C**), indicating the Ag85 activity is dispensable for the formation of Ag85+ storage compartments and bacterial replication in human macrophages. To confirm these findings, we generated a Mtb strain knocked out for Ag85B (Mtb *ΔfbpB)* (**Supp. Figure 6A and 6B**) to test if loss of Ag85B had an impact on Ag85+ MASC and bacterial replication. By comparing macrophages infected with Mtb WT and Mtb *ΔfbpB,* we observed that loss of Ag85B has negligible effect on the number and morphology of Ag85+ MASC and bacterial replication (**Figure 4D to 5F**). Together, our findings indicate that whereas Ag85B accumulates in the MASC, it does not functionally contribute to their formation in human macrophages.

**Figure 6.**
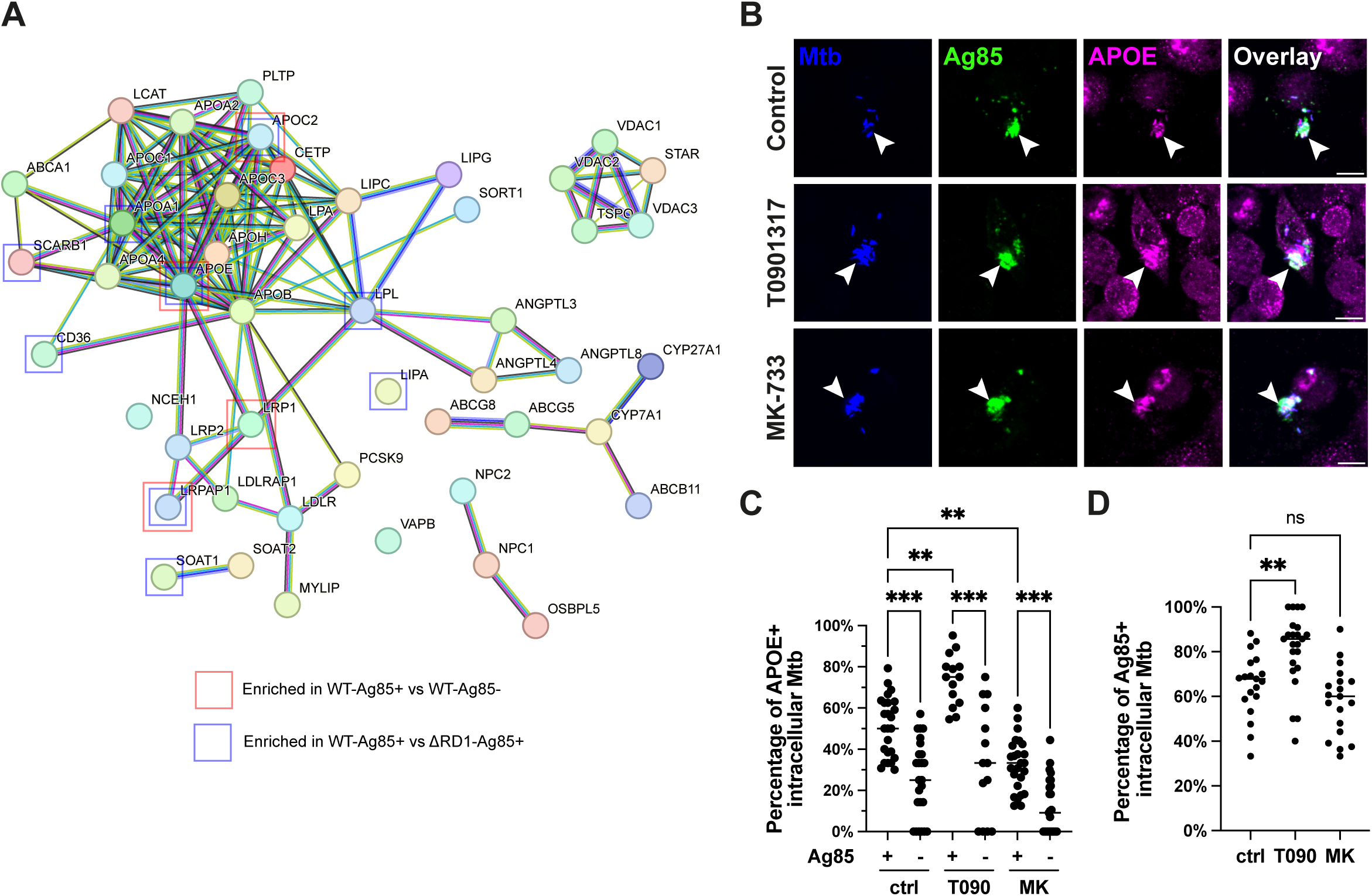
The Ag85+ antigen storage compartment is active in cholesterol metabolism and recruits the lipid binding protein APOE. (A) STING analysis of human proteins in cholesterol metabolism of KEGG pathway analysis. Red and blue squares depict proteins enriched in Ag85+ structures relative to Ag85- Mtb WT and Ag85+ structures induced by Mtb WT relative to ΔRD1, respectively. (B) Representative confocal images of control, T0901317 and MK-733-treated THP-1 cells infected by Mtb WT at 3h pi, images showing Ag85 (green), APOE (magenta) and Mtb (blue). Scalebar = 10 µm. (C) Quantification of Ag85+ Mtb in control, T0901317 (T090) and MK-733 (MK)-treated THP-1 cells infected by Mtb WT at 3h pi. (D) Quantification of percentage of APOE+ in Ag85+ and Ag85- Mtb in control, T0901317 (T090) and MK-733 (MK)-treated THP-1 cells infected by Mtb WT at 3h pi.

### The Ag85+ storage compartment is remodeled by the Mtb ESX-1 secretion system

To further understand the functional role and composition of MASC, we then determined the protein composition of the Ag85+ MASC using an spatial proteomics approach^36^. THP-1 cells infected with Mtb WT and Mtb ΔRD1 were stained for Ag85 and imaged on a fluorescence microscope. The captured Ag85 and Mtb channels were thresholded and segmented to generate the masks of Ag85 and Mtb, respectively. Ag85+ and Ag85- Mtb regions of interest (ROIs) were generated and illuminated using a two-photon laser source to photo-biotinylate the ROIs, where the biotinylated cells were further dissociated, enriched for biotinylated proteins, digested and identified using mass spectrometry (**Figure 5A and Supplementary Table 1**). As a control for the specificity of the photo-biotinylation step, we detected a significant enrichment of Ag85A, Ag85B and Ag85C in Ag85+ samples as compared to Ag85- samples (**Supp. Figure 7A**).

We detected a striking enrichment of Mtb proteins in the Ag85⁺ storage compartment induced by Mtb WT, confirming that the MASC selectively confines secreted Mtb proteins (**Figure 5B**). There was a significant enrichment of carbon, amino acid and fatty acid metabolic pathways as well as structural and secreted factors of the ESX-1 and ESX-3 secretion systems, which also include previously reported Mtb antigens presented by human macrophages^37^ (**Figure 5C and Supp. Figure 7B to 7D**). We then performed KEGG pathway analysis on the enriched macrophage proteins in the Ag85+ storage compartment induced by Mtb WT (**Figure 5D**). We observed a strong enrichment of endocytic trafficking and vesicular trafficking proteins, in particular SNAREs as well as metabolic pathways (**Figure 5E and Supp. Figure 8A**). Interestingly, we observed a strong enrichment of glycan degradation pathways (**Figure 5E**).

To identify the ESX-1 dependent changes in the Ag85+ MASC, we compared the enrichment of proteins in the Ag85+ MASC induced by Mtb WT and Mtb ΔRD1 (**Figure 5F**). Among the Mtb proteins enriched in Ag85+ storage compartment induced by Mtb WT in an ESX-1 dependent manner, we found metabolic and biosynthetic pathways, indicating an ESX-1- dependent remodeling of Ag85+ MASC and potential Mtb metabolic adaptation to distinct microenvironment (**Figure 5G**). Furthermore, we detected an enrichment of Mtb antigens (**Supp. Figure 9A**), enrichment of ESX-1 components (**Supp. Figure 9B and 9C**) and significant depletion of ESX-3 components (**Supp. Figure 9D and 9E**) in Mtb WT induced Ag85+ structures. The ESX-1 dependent differential protein enrichment was also reflected in the macrophage components of the Ag85+ MASC (**Figure 5H**). Moreover, we observed an enrichment in glycan degradation proteins as well as lysosome and cholesterol metabolism pathways in Ag85+ MASC induced by Mtb WT (**Figure 5I and Supp. Figure 10A**). Together, these results indicate that the Ag85+ storage compartment undergoes a pronounced remodeling via the ESX-1 secretion system.

### The Ag85+ storage compartment is enriched in metabolic pathway proteins and modulated by Mtb ESX-1

Our analysis showed an enrichment of cholesterol metabolism pathways in Ag85+ compartments induced by Mtb WT such as apolipoprotein A1, C2 and E (**Figure 6A**). Apolipoproteins facilitate cholesterol transport, a pathway implicated in Mtb restriction in the context of general metabolic state^38,39^ and neutrophil extracellular traps^39^, but the role of apolipoproteins in macrophage-Mtb interactions is less known. We detected an enrichment of Apolipoprotein E (APOE) in Ag85+ compartments induced by Mtb WT and compared to Mtb ΔRD1, suggesting a potential ESX-1-dependent regulation of the metabolic activity of the Ag85+ MASC (**Figure 6A**). We confirmed the predominant localization of Apolipoprotein E (APOE) to Ag85+ MASC (**Figure 6B and 6C)**. To determine if the incidence of Ag85+ structures or its colocalization with APOE will be altered by the cellular cholesterol level and export, we treated the macrophages with Liver X Receptor (LXR) agonist, T0901317 that elevates *APOE* expression and cholesterol export^40^ and HMG-CoA reductase inhibitors, simvastatin (MK-733) that reduces *APOE* expression^41^. We observed an elevation in Ag85+ MASC incidence upon T0901317 treatment, while MK-733 treatment has negligible effect (**Figure 6D**). The pharmacological perturbation of cellular cholesterol metabolism does not affect the preferential APOE localization to Ag85+ storage compartment, while treatment of T0901317 promotes the localization of APOE to Ag85+ storage compartment, and MK-733 exerts opposite effects (**Figure 6C**). Altogether, we have validated the preferential localization of APOE to Ag85+ MASC and a cholesterol metabolism-driven modulation on compartment composition triggered by Mtb in human macrophages.

## Discussion

Understanding when and where Mtb antigens are secreted in infected antigen presenting cells is important for the development of protective TB vaccines. During infection, Mtb dwells within a phagosome upon phagocytosis by macrophages, where the secreted bacterial components are localized in phagosomes and endocytic compartments^7,8,11,42^.

Here, we report the development of a split fluorescent reporter system to study antigen secretion and localization in infected cells. As only complementation in the cytosol will result in florescence signal, this system allows to spatiotemporally track the translocation of antigens into the cytosol and between cells. As proof of concept, we used Ag85B, one of the most relevant TB vaccine antigen leads, and showed that ESX-1 dependent phagosome damage is an important step in the access of this antigen to the cytosol. Using this system, we also discovered that Ag85B can be associated with macrophage membranes with potential consequences for antigen processing. Importantly, this reporter can be adapted and expanded to virtually study any Mtb secreted antigen in infected cells.

By combining different imaging modalities with the Ag85B reporter and endogenous Ag85 localization, we propose here that Mtb trigger two types of Mtb antigen storage compartments (MASC). The MASC we report here are large host endocytic compartments proximal to Mtb, being distinct from the previous reported AEV^29^. Of the two types of MACS, the first one is an intact MASC where Ag85 accumulates and from where Ag85 loaded vesicles potentially originate from the phagosomes. These vesicles are transported to plasma membrane and transfer to other cells as described before ^6,7,29,43^. The second type of MASC, with damaged membranes enabled by the Mtb ESX-1 secretion system, permits Ag85B not only to localize within the compartment but also to access the cytosol, associate with phagosome membranes, and accumulate in vesicles that can be transferred to other cells. Our CLEM data shows that it is likely that this MASC is leaky and composed of many membranes. This MASC is related to the tubulovesicular compartments that are triggered by the Mtb ESX-1 secretion system in macrophages^22–24^. The existence of two distinct compartments raises some important cell biology and immunology questions. It is likely that in the intact MASC, antigens are processed and presented via the MHC Class II to the cell surface. In contrast, because of its leakiness, the damaged MASC is likely to play a role in both MHC Class I and II antigen processing and loading. In this context, our finding of two types of MASC, one that is intact where antigens are processed by lysosomal activity and a second damaged MASC where antigens access the cytosol, conciliate findings showing that T7SS substrates are a prominent source of Mtb peptides presented on MHC-I^25^ and mycobacterial peptides identified included twin arginine translocation (Tat) pathway substates, Sec pathway secretion substrates, membrane-associated proteins, and cytosolic proteins^37^. We observed 24 of our enriched Mtb proteins among the reported presented antigens, including but not limited to Ag85A, Ag85B, Ag85C, mmpL3, embA, embB and fadD26^37^. Future work will define how the balance and time of formation of these two compartments impact antigen presentation. The dispensability of Ag85B activity suggest that Ag85 is a payload instead of a driver to the formed membrane compartments and it is likely other Mtb antigens are also sequestered in this compartment.

The distinct morphological features of the intact vs. damaged MASC suggest a differential composition and potential different physiological function. The spatial proteomics study delineated some of the components that contribute to these differences. Our data indicate that Mtb in the MASC are enriched in metabolic pathway components as the main differences between Ag85+ MASC and Ag85- Mtb were the significant enrichment in several metabolic pathways linked to lipid metabolism. We also found that bacteria in the Ag85+ MASC are enriched in the ESX-3 secretion system, suggesting that bacteria respond to iron and zinc in the MASC or stress in general^44^. Interestingly, the components of the ESX-3 secretion system are classified as CD4^+^ T cell antigens^45^. Notably, the MASC is enriched in cholesterol and purine metabolic pathway components. Moreover, the MASC is enriched in lysosomal components and lysosomal function, suggesting MASC could harbor degradative functions. In agreement with the notion that the MASC constitutes of damaged membranes, there was a significant enrichment of the ESCRT machinery components.

Our data also highlight that Mtb not only trigger the formation of a damaged MASC via the ESX-1 but also that the activity of ESX-1 induces a remarkable metabolic remodeling of the MASC. We observed an enrichment of metabolic pathways, in particular cholesterol metabolism in MASC, consistent with the association of Mtb to lipid droplets for replication^46–48^. In agreement with an increase in cholesterol metabolism, we found that APOE is recruited to the MASC and the perturbation of cholesterol metabolism alters the recruitment of APOE to MASC. A role of APOE in mycobacterial lipid antigen presentation has been reported^13^ and we provide evidence that this could be linked to the formation of MASC enriched in distinct metabolic pathway components. Further studies to define a role for APOE regulation of cholesterol metabolism and MASC shall shed light on the molecular mechanism underlying the regulation of MASC composition and its implication to Mtb survival. Nonetheless, we discovered the MASC contains Ag85 as a payload and this suggest an unprecedented sorting mechanism of antigens than give rise to different subtypes of MASC. Future work shall reveal the heterogeneity of MASC as well as their functions in various antigen presenting cell types.

Altogether, using a novel reporter approach combined with subcellular spatial proteomics, we report here the identification of two distinct ESX-1 dependent Mtb antigen storage compartments that are rapidly formed after Mtb phagocytosis. These compartments are plausibly important in nutrient acquisition by Mtb as well as antigen presentation by macrophages and the regulation of T cell responses. Understanding how Mtb antigens are trafficked and processed could help to identify TB protective vaccine targets and strategies.

## Acknowledgements

We thank the Advanced Light Microscopy, Flow Cytometry and Proteomics facilities at the Crick for their support in various aspects of the work. This work was supported by the Francis Crick Institute (to M.G.G.), which receives its core funding from Cancer Research UK (CC2081), the UK Medical Research Council (CC2081), and the Wellcome Trust (CC2081). For the purpose of Open Access, the author has applied a CC BY public copyright license to any Author Accepted Manuscript version arising from this submission.

## Author Contributions

Conceptualization: CHL., MGG.; Methodology: CHL., TA., AF., CC.; Software: TA.; Validation: CHL.; Formal Analysis: CHL., TA.; Investigation: CHL., TA., TF.; Resources: CHL., JOC., PKL.; Data curation: CHL., TA.; Writing original draft: CHL., MGG.; Writing review & editing: all authors.; Visualization: CHL.; Supervision: MGG., CHL.; Project administration: MGG., CHL.; Funding acquisition: MGG.

## Supplementary figures

Supplementary Figure 1. Characterization of Ag85B split reporter by fluorescence microscopy. (A) Representative confocal images of Mtb H37Rv WT and WT Ag85B-7xmNG_11_ infected THP-1 WT and THP-1 mNG_1-10_ at 3h and 72h pi, images showing nuclei (blue), mNG (green) and Mtb (magenta). Scalebar = 20 µm. (B) Quantification of mNG+ structures in THP-1 mNG_1- 10_ and THP-1 WT cells infected by Mtb H37Rv WT and WT Ag85B-7xmNG_11_ at 3h and 72h pi.

Supplementary Figure 2. Characterization of Ag85B split reporter by flow cytometry. (A) Flow cytometry plots showing the gating strategy of the analysis of THP-1 mNG_1-10_ cells. (B) Flow cytometry plots showing the Mtb-infected cells of the analysis of Ag85B split reporter.

Supplementary Figure 3. Validation of the binding specific of Ag85B antibody (RRID:AB_776575) to Ag85 proteins. (A) Representative confocal images of THP-1 cells infected by Mtb H37Rv WT and *ΔfbpB* at 3h pi, images showing nuclei (blue), Ag85 (green) and Mtb (magenta). Scalebar = 10 µm. (B) Pairwise comparison of enriched and depleted proteins bound by Ag85B antibody (RRID:AB_776575) in immunoprecipitation as compared to control antibody. Plots showing –log_10_(p value) from two-sided Student’s t test and log_2_FC differences between each comparison.

Supplementary Figure 4. Ag85+ structures lack antigen export vesicles marker (A) Representative confocal images of THP-1 cells infected by Mtb H37Rv WT and ΔRD1 at 3h and 24h pi, images showing Ag85 (green), Tubulin (magenta) and Mtb (blue). Scalebar = 10 µm. (B) Representative scatter plot of Tubulin intensity against Ag85 intensity of Mtb WT and ΔRD1 in THP-1 cells at 3h and 24h pi.

Supplementary Figure 5. Complemented Ag85B split reporter transfers from cell to cell (A) Schematic of labelling mechanism of Mtb Ag85B split fluorescent reporter and antibody staining to monitor antigen localization in bystander macrophages. (B) Quantification of mNG+ uninfected THP-1 mNG_1-10_ cells among cells infected by Mtb H37Rv WT and ΔRD1-Ag85B-7xmNG_11_ at 3h and 72h pi. (C) Schematic of cell-to-cell transfer of cytosolic Ag85B-7xmNG_11_. (D) Representative confocal images of uninfected THP-1 cells among cells infected by Mtb H37Rv WT and ΔRD1 at 3h, 24h, 48h and 72h pi, images showing nuclei (blue), Ag85 (green) and Mtb (magenta). Scalebar = 10 µm. (E) Quantification of Ag85+ uninfected THP- 1 cells among cells infected by Mtb H37Rv WT and ΔRD1 at 3h, 24h, 48h and 72h pi.

Supplementary Figure 6. Characterization of Mtb *ΔfbpB* mutant (A) Genotyping of Mtb H37Rv WT and *ΔfbpB* mutant by PCR showing amplicon of *fbpB* and h*sp60* promoter (positive control). Expected size of *fbpB* amplicon is depicted by *. (B) Western blot probing Ag85 and GroEL of Mtb H37Rv WT and *ΔfbpB* mutant. Expected size of Ag85B and Ag85B-7xmNG_11_ are depicted by * and #, respectively.

Supplementary Figure 7. Enriched Mtb components in Ag85+ structures induced by Mtb WT. (A) Pairwise comparison of enriched and depleted Mtb proteins in Ag85+ structures relative to Ag85- Mtb WT. White dots depict Ag85A, Ag85B and Ag85C. (B) Pairwise comparison of enriched and depleted Mtb proteins in Ag85+ structures relative to Ag85- Mtb WT. White dots depict detected ESX-1 components. (C) Pairwise comparison of enriched and depleted Mtb proteins in Ag85+ structures relative to Ag85- Mtb WT. White dots depict detected ESX-3 components. (D) STRING analysis of enriched Mtb components in Ag85+ structures relative to Ag85- Mtb WT. Enriched proteins of KEGG pathways “Metabolic Pathways”, “Ribosome”, “Arabinogalactan biosynthesis - Mycobacterium” and “Bacterial secretion system” are colored in red, blue, yellow and green, respectively. Protein clusters of “Ag85 complex”, “ESX-1 secretion system” and “ESX-3 secretion system” are highlighted. Plots showing -log_10_(p value) from two-sided Student’s t test and log_2_FC differences between each comparison.

Supplementary Figure 8. Enriched host components in Ag85+ structures induced by Mtb WT. (A) STRING analysis of enriched host components in Ag85+ structures relative to Ag85- Mtb WT. Enriched proteins of KEGG pathways “SNARE interactions in vesicular transport”, “Lysosome” and “Metabolic pathways” are colored in red, blue and green, respectively. Protein clusters of “ESCRT complex”, “Cholesterol metabolism”, “Translation” and “Nuclear protein-containing complex” are highlighted.

Supplementary Figure 9. Enriched and depleted Mtb components in Ag85+ structures induced by Mtb WT relative to ΔRD1 (A) Pairwise comparison of enriched and depleted Mtb proteins in Ag85+ structures induced by Mtb WT relative to ΔRD1. White dots depict Ag85A, Ag85B, Ag85C and mpt64. (B) Pairwise comparison of enriched and depleted Mtb proteins in Ag85+ structures induced by Mtb WT relative to ΔRD1. White dots depict detected ESX-1 components. (C) STRING analysis of enriched Mtb proteins in Ag85+ structures induced by Mtb WT relative to ΔRD1. Enriched proteins of KEGG pathways “Metabolic Pathways”, “Ribosome”, “Arabinogalactan biosynthesis – Mycobacterium” and “Fatty acid metabolism” are colored in red, blue, yellow and green, respectively. Protein clusters of “Ag85 complex” and “ESX-1 secretion system” are highlighted. (D) Pairwise comparison of enriched and depleted Mtb proteins in Ag85+ structures induced by Mtb WT relative to ΔRD1. White dots depict detected ESX-3 components. (E) STRING analysis of depleted Mtb proteins in Ag85+ structures induced by Mtb WT relative to ΔRD1. Protein cluster of “ESX-3 secretion system” is highlighted. Plots showing -log_10_(p value) from two-sided Student’s t test and log_2_FC differences between each comparison.

Supplementary Figure 10. Enriched host components in Ag85+ structures induced by Mtb WT relative to ΔRD1. (A) STRING analysis of enriched host components in Ag85+ structures induced by Mtb WT relative to ΔRD1. Enriched proteins of KEGG pathways “Phagosome”, “Lysosome” and “Metabolic pathways” are colored in red, blue and green, respectively. Protein clusters of “ESCRT complex”, “Cholesterol metabolism” and “Ribonucleoprotein complex” are highlighted.

**Movie S1: Dynamic association of Ag85B-7xmNG11 with Mtb taking place from 2 hours after infection, Related to Figure 3B**.

**Movie S2: Dynamic association of Ag85B-7xmNG11 with Mtb taking place from 2 hours after infection, Related to Figure 3B**.

**Movie S3: Dynamic association of Ag85B-7xmNG11 with Mtb taking place from 24 hours after infection, Related to Figure 3C**.

## Materials and Methods Cell culture

THP-1 (TIB-202) human monocyte cell line was obtained from American Type Culture Collection (ATCC) and HEK-293FT human embryonal kidney cells was obtained from ThermoFisher Scientific and were used within 20 passages of receipt. THP-1 cells were cultured in RPMI-1640 medium (ThermoFisher Scientific) containing 10% (v/v) heat-inactivated fetal bovine serum (FBS, ThermoFisher Scientific) at 37°C with 5% CO_2_ and 100% humidity. THP-1 cells were differentiated in RMPI-1640 + 10% FBS + 10 µg/ml phorbol 12-myristate 13-acetate (PMA, Merck) for 24 hours and rested in RMPI-1640 + 10% FBS prior to Mtb infection. HEK-293FT cells were cultured in Dulbecco’s Modified Eagle Medium (DMEM, high glucose, GlutaMAX Supplement, ThermoFisher Scientific) containing 10% FBS at 37°C with 5% CO_2_ and 100% humidity.

## Plasmid construction of split fluorescent reporter

Plasmids harboring mNeonGreen3K (human codon optimized) and 7xmNeonGreen3K(11) gene fragments were synthesized and purchased from GENEWIZ Germany GmbH. The DNA sequence of mNG3K(1-10) was amplified using PCR and inserted into pLJM1 eGFP (Addgene, #19319) between AgeI and EcoRI sites to generate pLJM1-mNG3K(1-10). The DNA sequence of *fbpB* was amplified from genomic DNA of Mtb H37Rv and inserted upstream of the 7xmNG3K(11) sequence in a peGFP-N1 backbone. The *fbpB*-7xmNG3K(11) sequence was subsequently cloned downstream of the putative *fbpB* promoter sequence together with Psmyc-E2-Crimson^49^ into a replicating *Mycobacterium* plasmid backbone with hygromycin resistance marker using Gateway cloning to generate pMEH P*fbpB*-Ag85B-7xmNG_11_. The Plasmid DNA sequences are available upon request.

## Engineering of split fluorescent reporter in THP-1

To produce lentivirus harboring the mNG3K(1-10), HEK-293FT were transiently co-transfected using calcium phosphate with: (i) pLJM1-mNG3K(1-10) (construct plasmid), (ii) pCMV-VSV-G (envelop plasmid, Addgene) and (iii) pCMV-dR8.2 dvpr (packaging plasmid, Addgene). The produced lentivirus was harvested from the media and concentrated using Lenti-X™ Concentrator (Takara Bio) to achieve a 50-fold concentration following manufacturers protocol. THP-1 cells were transduced with produced lentivirus at MOI of 0.1 with 10 µg/ml polybrene (Merck) and being selected with 1 µg/ml puromycin (Invivogen) 3 days post-transduction. Transduced THP-1 cells were passaged twice in the presence of 1 µg/ml puromycin prior to single cell cloning using limited dilution. Single cell clones were cultured and validated for the insertion of mNG3K(1-10) cassette.

## Mycobacterium strains and culture conditions

Mtb H37Rv (WT) was a generous gift by Prof. Douglas Young (The Francis Crick Institute) and ΔRD1 was provided by Dr. Suzie Hingley-Wilson (University of Surrey). Mtb HN878 was obtained from BEI Resources (NR-13647). Mtb deletion mutant for *fbpB* (rv1886c) was constructed using ORBIT^50^. All Mtb mutants were validated by PCR and confirmed by whole genome sequencing. Mtb constitutively expressing E2-Crimson was generated by transformation with pTEC19 (Addgene, 30178, deposited by L. Ramakrishnan). Mtb harboring the expression cassette of Ag85B-7xmNG_11_ was generated by transformation with pMEH P*fbpB*-Ag85B-7xmNG ^51^. All Mtb strains were tested for phthiocerol dimycocerosate (PDIMs) positivity by thin-layer chromatography of lipid extracts. Mtb strains were cultured in Middlebrook 7H9 (Sigma-Aldrich, M0178) supplemented with 0.2% glycerol (Fisher Chemical, G/0650/17), 0.05% Tween-80 (Sigma-Aldrich, P1754) and 10% ADC (BD Biosciences, 212352) supplemented with appropriate selection markers 50 µg/ml Hygromycin B (Invitrogen, 10687010) or 25 µg/ml Zeocin (Invivogen, ant-zn-05).

## Mtb infection

Mtb cryovials were thawed and cultured in Middlebrook 7H9 supplemented with 0.2% glycerol, 0.05% Tween-80, 10% ADC and 50 µg/ml Hygromycin B until late log phase. The Mtb culture was collected by centrifugation at 3000 x g for 5 minutes, washed twice with PBS and dislodged by vortexing with glass beads for 1 minute. The dislodged bacteria pellets were resuspended in appropriate cell culture media and centrifuged at 1200 x g for 5 minutes to pellet bacterial clumps. The supernatant containing single bacterium were collected and measured with a spectrophotometer at OD600 to determine the bacterial concentration. The single bacterial suspension was diluted in appropriate cell culture medium to achieve a multiplicity of infection (MOI) of 1, unless otherwise specified. Cells were washed once with PBS and infected with the diluted bacterial suspension for 2 hours at 37 °C. After 2 hours of infection, cells were washed 3 times with PBS and cultured in appropriate medium at 37 °C (this is defined as time zero of the infection). At designated experimental endpoint, cells were washed once with PBS and fixed overnight in 4% PFA (diluted from 16% stock, Electron Microscopy Sciences) in PBS at 4 °C. The single bacterial suspension was pelleted by centrifugation at 3000 x g for 5 minutes and fixed overnight in 4% PFA in PBS at 4 °C.

## Immunofluorescence staining and microscopy

Cells were washed once with PBS and fixed in 4% PFA in PBS for 15 minutes at room temperature. PFA-fixed cells were washed three times with PBS and quenched in 50 mM NH_4_Cl (Merck) in PBS for 15 minutes. Cells were then washed once with PBS, permeabilized with 0.01% saponin (Merck) in 1% BSA (Cell Signaling Technology) PBS and blocked in 3% BSA PBS at room temperature for 1 hour each. Blocked cells were washed 3 times with PBS and incubated with primary antibodies for 1 hour at room temperature. The stained cells were washed 3 times with PBS and incubated with secondary antibodies for 1 hour at room temperature. The stained cells were washed 3 times with PBS and stained with 4’,6-diamidino-2-phenylindole (DAPI, Invitrogen) for 10 minutes at room temperature. The cells were then washed 3 times with PBS and mounted with Dako Omnis Fluorescence Mounting Medium (Agilent), dried overnight at room temperature and stored at 4°C.

The stained samples were imaged using Olympus CSU-W1 SoRa Spinning Disk Microscope (Olympus) using 20x 0.7 NA air, 40x 1.25 NA silicone, 60x 1.3 NA silicone and 60x 1.35 NA oil immersion objectives or STELLARIS 5 Confocal Microscope Platform (Leica Microsystems) using 40x 1.3 NA oil immersion objective.

## Flow cytometry

Mtb-infected cells were detached using TrypLE (Invitrogen) with cell scraping and fixed in overnight in 4% PFA in PBS at 4 °C. The fixed cells were washed once with PBS and quenched in 50 mM NH_4_Cl in PBS for 15 minutes. Quenched cells were washed once in PBS, resuspended in 1% BSA PBS and stored 4°C. PFA-fixed single bacterium suspensions were washed with PBS once and incubated with primary antibodies for 1 hour at room temperature. The stained bacteria were washed once with PBS and incubated with secondary antibodies for 1 hour at room temperature. The stained bacteria were washed once with PBS, resuspended in 1% BSA PBS and stored 4°C. Cells and Mtb in suspension were analyzed using a BD LSRFortessa™ Cell Analyzer (BD Biosciences) and analyzed using FlowJo software (FlowJo, LLC, v10.8.1). At least 10,000 events per condition were recorded.

## Treatment with chemical compounds

Ebselen (Tocris, 5245) was supplemented to Mtb culture in Middlebrook 7H9 medium 24 hours prior to Mtb infection and cell culture medium throughout the infection at 16 µM.

## Live correlative light electron microscopy

THP-1 cells were seeded on 35 mm gridded-glass bottom dish (Mattek, P35G-1.5-14-C-GRD) at 1x10^6^ cells per dish and infected with Mtb as described above. At 0 h post-infection, the dishes were sealed with parafilm, mounted onto a dish holder and loaded onto STELLARIS 5 Confocal Microscope Platform (Leica Microsystems) equipped with an environmental chamber (Okolab) set at 37°C with 5% CO_2_ to acclimatize for 1 hour prior to imaging using a 40x 1.3 NA oil immersion objective.

Samples were fixed by adding a mixture of 8% PFA in 200 mM HEPES (pH 7.4) buffer to culture medium (v/v) and incubated at room temperature for 15 minutes and then replaced with 4% PFA in 100 mM HEPES overnight at 4°C before imaging by confocal microscopy. After staining with DAPI in PBS for 20 minutes, samples were first imaged by a laser scanning confocal microscope (Leica STELLARIS 5) and then changed to 1% glutaraldehyde in 200 mM HEPES (pH 7.4) for 30 minutes at room temperature. Fluorescently imaged samples were processed for CLEM in a Biowave Pro (Pelco, USA) with use of microwave energy and vacuum. Cells were twice washed in HEPES (Sigma-Aldrich, H0887) at 250 W for 40 s and postfixed using a mixture of 2% osmium tetroxide (Taab O011) and 1.5% potassium ferricyanide (Taab, P018) (v/v) at an equal ratio for 14 minutes at 100-W power [with/without vacuum, 20 inches (67,727.8 Pa) Hg at 2-minutes intervals]. Samples were washed with distilled water twice on the bench and twice again in the Biowave 250 W for 40 s. Samples were stained with 1% aqueous uranyl acetate (Agar Scientific AGR1260A) in distilled water (w/v) for 14 minutes at 100-W power [with/without vacuum, 20 inches (67,727.8 Pa) Hg at 2- minutes intervals] and then washed using the same settings as before. Samples were dehydrated using a stepwise acetone series of 50, 75, 90, and 100% and then washed 4x in absolute acetone at 250 W for 40 s per step. Samples were infiltrated with a dilution series of 25, 50, 75, and 100% Ultra Bed Low Viscosity Epoxy Kit (EMS, 14310) (v/v) resin to propylene oxide. Each step was for 3 minutes at 250-W power [with/without vacuum, 20 inches (67,727.8 Pa) Hg at 30-s intervals]. Samples were then cured for a minimum of 48 hours at 60°C.

## Sample trimming and Scanning Electron Microscopy array tomography acquisition

Glass cover slips were removed by plunging samples into liquid nitrogen. Referring to grid coordinates, the sample block was trimmed, coarsely by a razor blade and then finely trimmed using a 35° ultrasonic, oscillating diamond knife (DiATOME, Switzerland) set at a cutting speed of 0.6 mm/s, a frequency set by automatic mode, and a voltage of 6.0 V, on an ultramicrotome EM UC7 (Leica Microsystems, Germany) to remove all excess resin surrounding the region of interest. Serial sections were collected on ITO (indium tin oxide) glass slides for imaging using a Zeiss Gemini SEM 460.

## CLEM image alignment

Fluorescence .LIF files were converted to .tif file format, and linear adjustments made to brightness and contrast using FIJI (version 2.14.0/1.54t). Fluorescence images were aligned to serialEM micrographs (TrakEM2) using the BigWarp_fiji_7.0.7 plugin. No less than 10 independent fiducials were chosen per alignment for 3D image registration. When the fiducial registration error was greater than the predicted registration error, a nonrigid transformation (a nonlinear transformation based on spline interpolation, after an initial rigid transformation) was applied.

## Image-based single bacterium analysis

To quantify the presence of Ag85+ compartments proximal to Mtb during infection, the image stacks were loaded into FIJI (version:2.9.0/1.54f) using Bio-Formats Importer, Z-projections were generated using maximum projection. Individual cell ROI was segmented by thresholding the cellular marker channel and converting the binary image mask to selection. Individual Mtb ROI was segmented by thresholding the E2-Crimson marker channel and converting the binary image mask to selection. Intracellular Mtb masks were generated by using the “AND” option of Image Calculator, which were then expanded using the “Dilate” function to capture Mtb proximity. The dilated Mtb masks were used to measure the mean fluorescence intensity of the cellular marker channels using “Analyze Particles”. The positivity of cellular markers was determined by the threshold value defined by the average fluorescent intensity of the dilated Mtb mask in control cells. The same workflow was used to determine the colocalization of markers proximal to intracellular Mtb.

## Immunoprecipitation using Ag85B antibody

THP-1 cells were infected with Mtb WT as described in the above section. At 3 hours after infection, uninfected and infected THP-1 cells were washed once with PBS and incubated in lysis buffer (Tris 8.0, 50 mM HEPES pH 8.0, 0.2% TritionX-100, 50 mM KCl) at room temperature for 5 minutes. The cells were then scrapped, and the cell lysates were collected, cell debris and nuclei were removed by centrifugation at 500 rcf for 10 minutes at 4°C. The cell lysates were then heated at 95°C for 25 minutes on a heat block prior to removal from biosafety level 3 laboratory. Dynabeads (ThermoFisher Scientific, 10765583) were crosslinked to Ag85B antibody (RRID:AB_776575) according to standard procedure. Cell lysates were incubated with the 20 µl of beads for 30 minutes at 4°C with constant rotation. Beads were washed with buffer containing 300 µM NaCl and 1% TritonX-100 to eliminate non-specific binding. Samples were eluted from Dynabeads using 2% SDS and digested using S-trap micro protocol according to manufacturer instructions (PROTIFI, C002-MICRO). 50 ng of peptide material was loaded onto individual Evotips per manufacturer instructions and subjected to LC MS/MS analysis. Peptides were resolved on Aurora Elite 15 cm column (Ion Optics) using Evosep One LC system using Whisper Zoom 40SPD method. For data acquisition TIMS TOF Ultra was operated in DIA-PASEF mode with the variable windows. Scan width was set between 100 and 1700 m/z, ion mobility window 1/K0 0.64 - 1/KO 1.45, ramp time - 150 ms and accumulation time -150 ms. Raw data was processed in Spectronaut 20 (Biognosys) against Uniprot human proteome (1 gene per protein) and Mycobacterium tuberculosis databases with standard settings. A log2(FC) of 0.58 was applied to determine enriched and depleted proteins.

## Spatial proteomics

THP-1 mcherry-LC3B cells were seeded in 2 Well chambered cover glass (Cellvis, C2-1.5H-N) at 1x10^6^ cells per chamber and infected with Mtb WT E2-Crimson or Mtb ΔRD1 E2- Crimson at MOI = 2 as described above. Infected cells were fixed overnight with 4% PFA at 3h and 24h post-infection. Fixed cells were stained with rabbit anti-Ag85 followed by AlexaFlour568 goat anti-rabbit following the staining protocol described above. The stained cells were then blocked with Block 1 and Block 2 buffers (SynCell, SYN-RI0106) according to manufacturer’s protocol. The masks of Ag85+ structures and Ag85- Mtb were defined as depicted in Figure 4a, the region of interest were then photolabelled using Labeling Reagent (SynCell) according to manufacturer’s protocol. The photolabeling reaction was quenched using Quench Reagent (SynCell) and stored at 4 °C until mass spectrometry sample preparation.

Following the labelling, cells were washed in water once, covered with 100 µl of scraping buffer (50 mM Tris-HCl pH = 8) and scraped from the cover slip. Scraped samples were sonicated for 5 minutes using BioRaptor nano (operated in 30 seconds on/30 seconds off mode) and treated with 50 U of Benzonase per sample for 30 minutes at RT. Samples were combined with 400 ul of enrichment buffer (50 mM Tris-HCl pH = 8, 0.01% DDM, 0.1% SDS) and incubated at 90 °C for an hour. Following 5 minutes of sonication, lysates were cleared at 20000 x g for 10 minutes. Supernatants were mixed with 30 µl of MagReSyn Streptavidin beads (ReSyn Biosciences), and samples were incubated for 1 hour with shaking. Beads were washed 5 times in washing buffer (100 mM TEAB, 0.01%DDM) and resuspended in 30 µl of 100 mM TEAB and 50 ng of Trypsin (Promega VA9000). Following overnight incubation, 50 ng of peptide material was loaded onto individual Evotips per manufacturer instructions and subjected to LC MS/MS analysis. Peptides were resolved on Aurora Elite 15 cm column (Ion Optics) using Evosep One LC system using Whisper Zoom 40SPD method. For data acquisition TIMS TOF Ultra was operated in DIA-PASEF mode with the variable windows. Scan width was set between 100 and 1700 m/z, ion mobility window 1/K0 0.64 - 1/KO 1.45, ramp time - 150 ms and accumulation time -150 ms. Raw data was processed in Spectronaut 20 (Biognosys) against Uniprot human proteome (1 gene per protein) and Mycobacterium tuberculosis databases with standard settings. A log2(FC) of 0.58 was applied to determine enriched and depleted proteins that were submitted to STRING pathway analysis^52^.

## Statistical analysis

Statistical analyses were performed using GraphPad Prism software Version 10.1.1 (270).

