## Supplementary Figures 1-10 for "ESX-1 remodels the *Mycobacterium tuberculosis* antigen storage compartment in macrophages"

SuppFigure 1

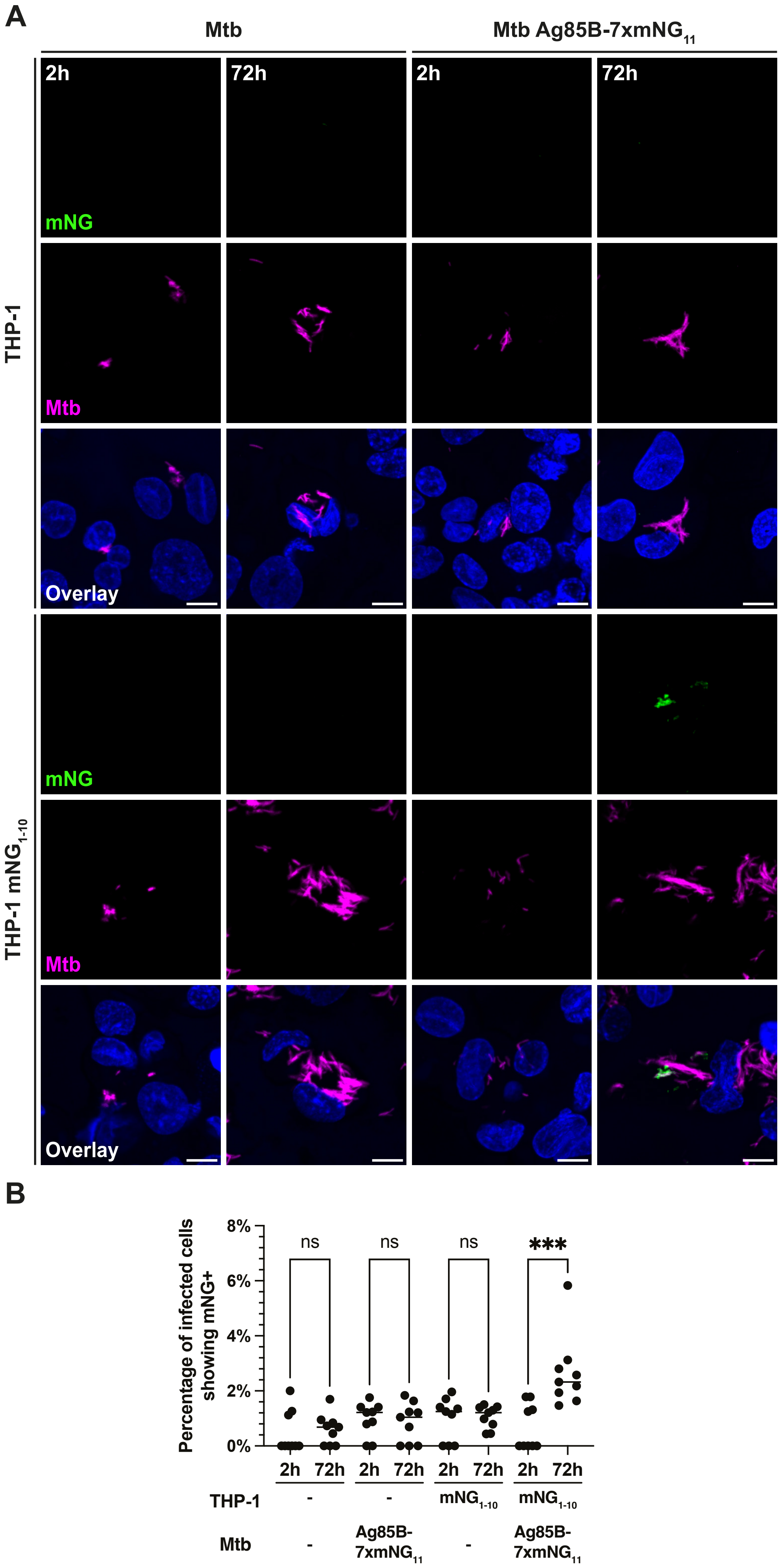

Supp. Figure 2

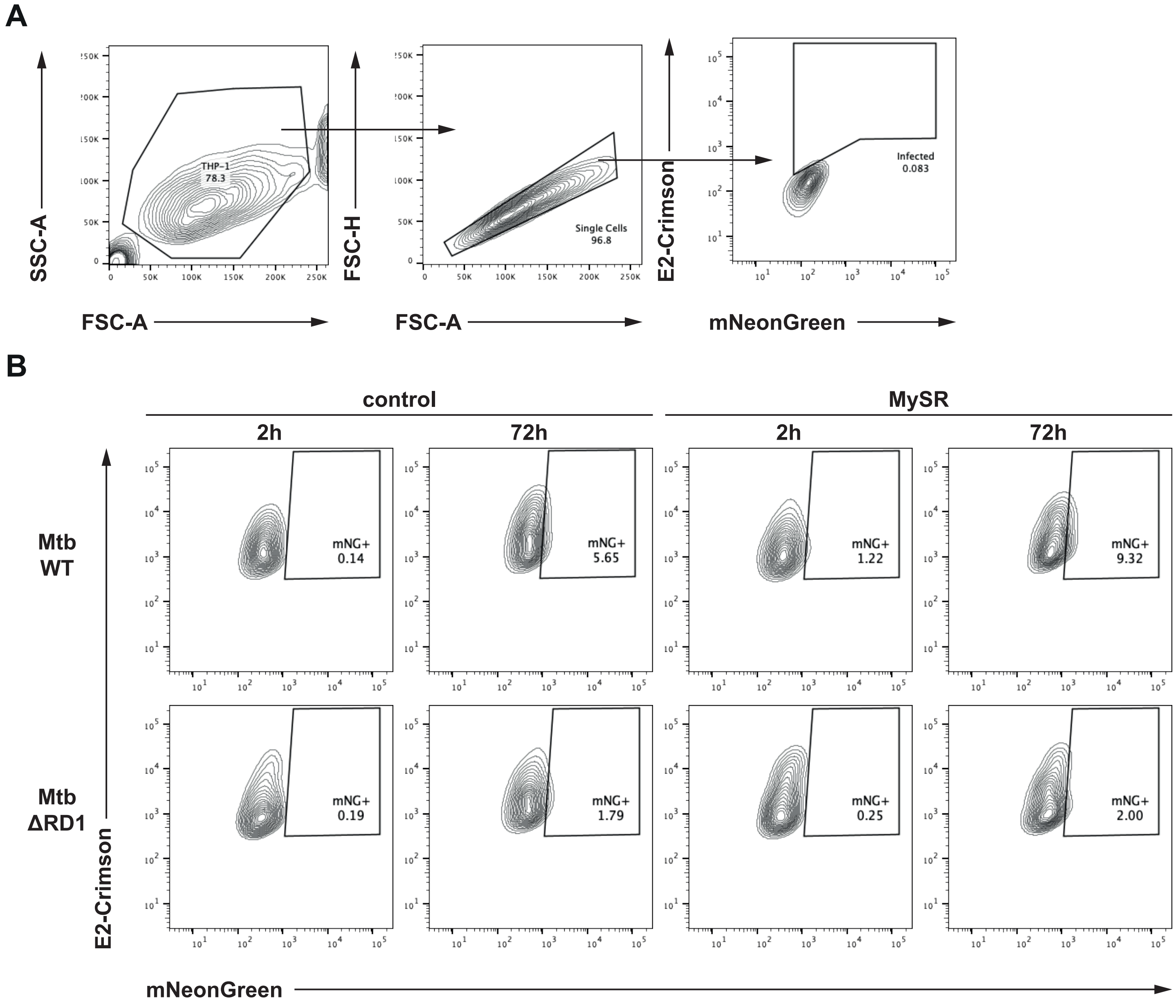

Supp. Figure 3

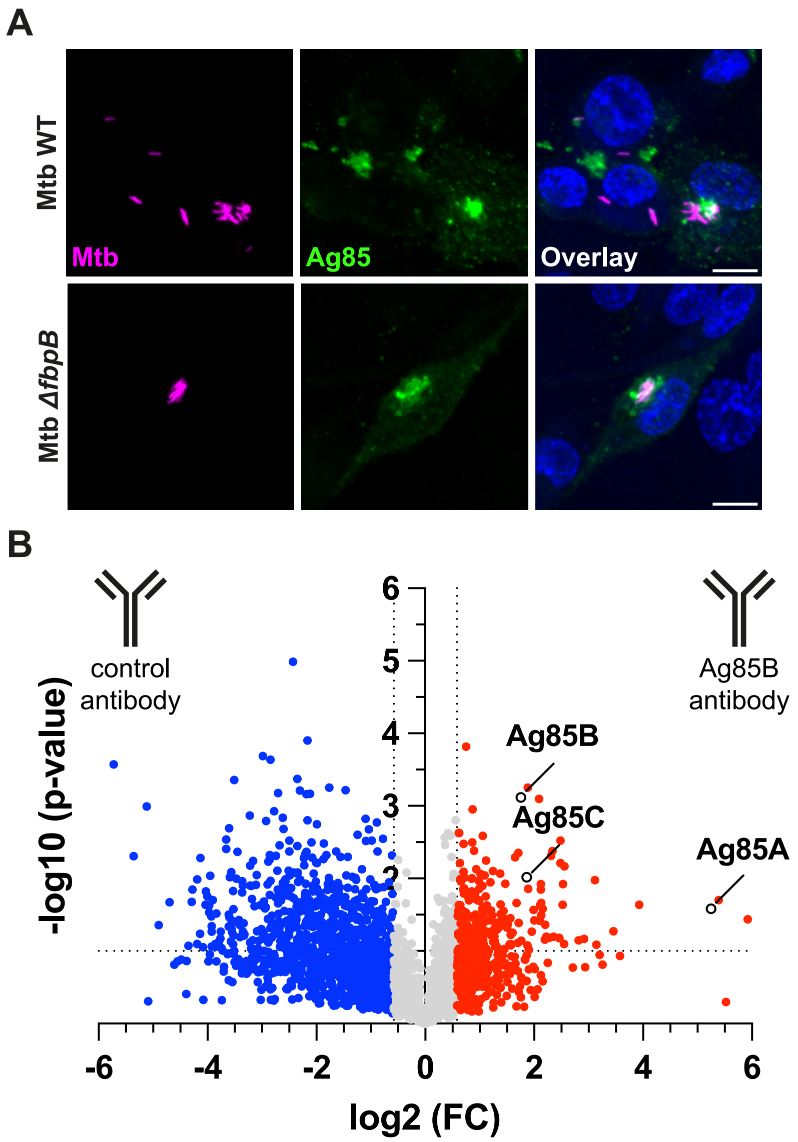

Supp. Figure 4

A

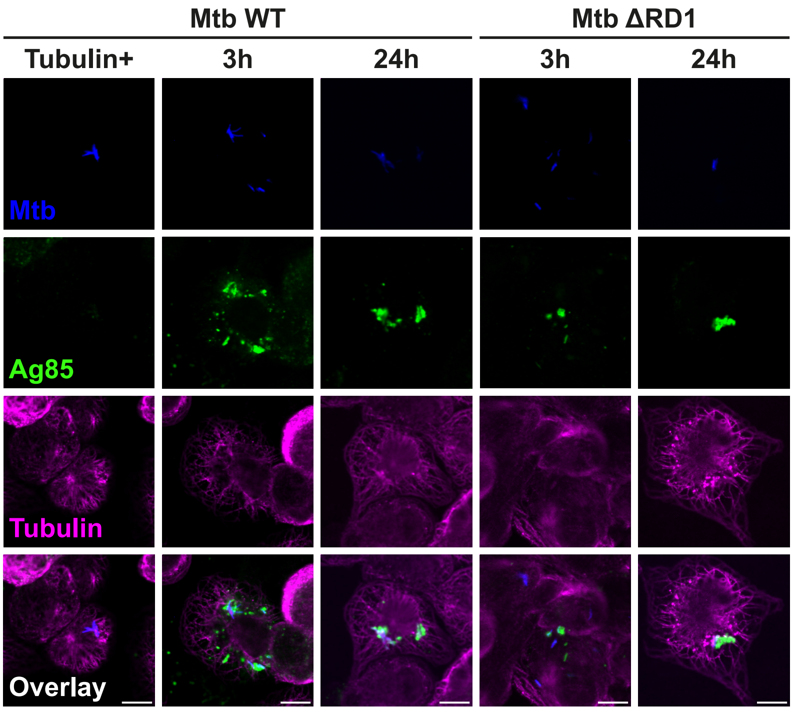


B

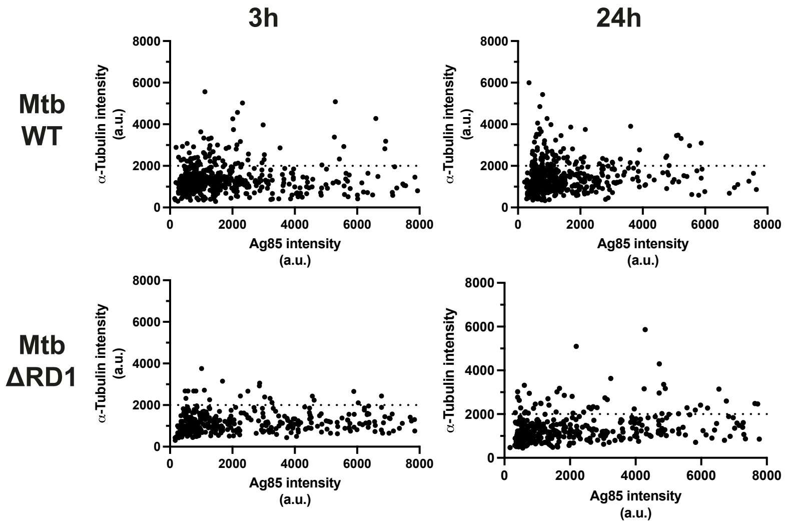

Supp. Figure 5

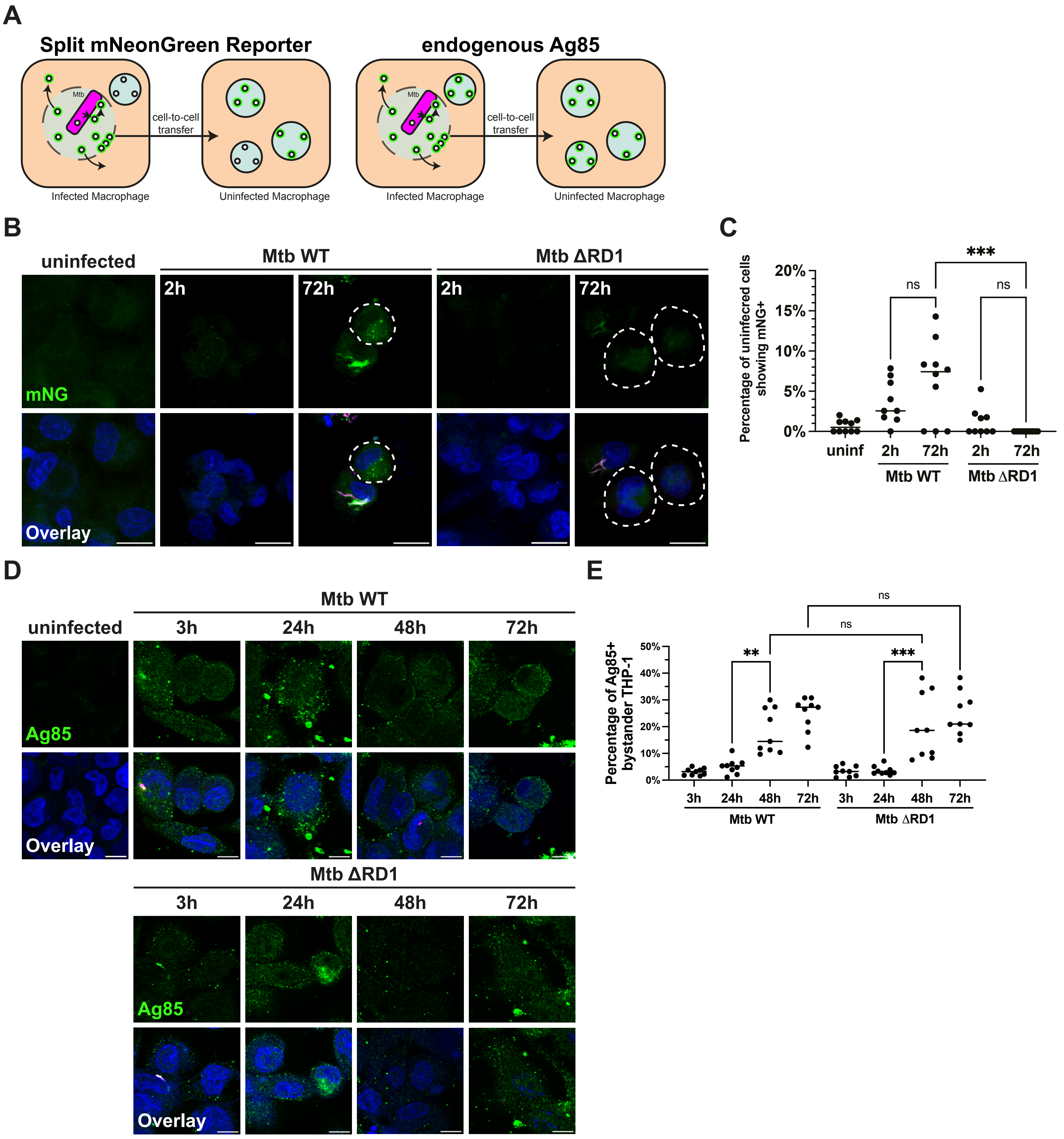

Supp. Figure 6

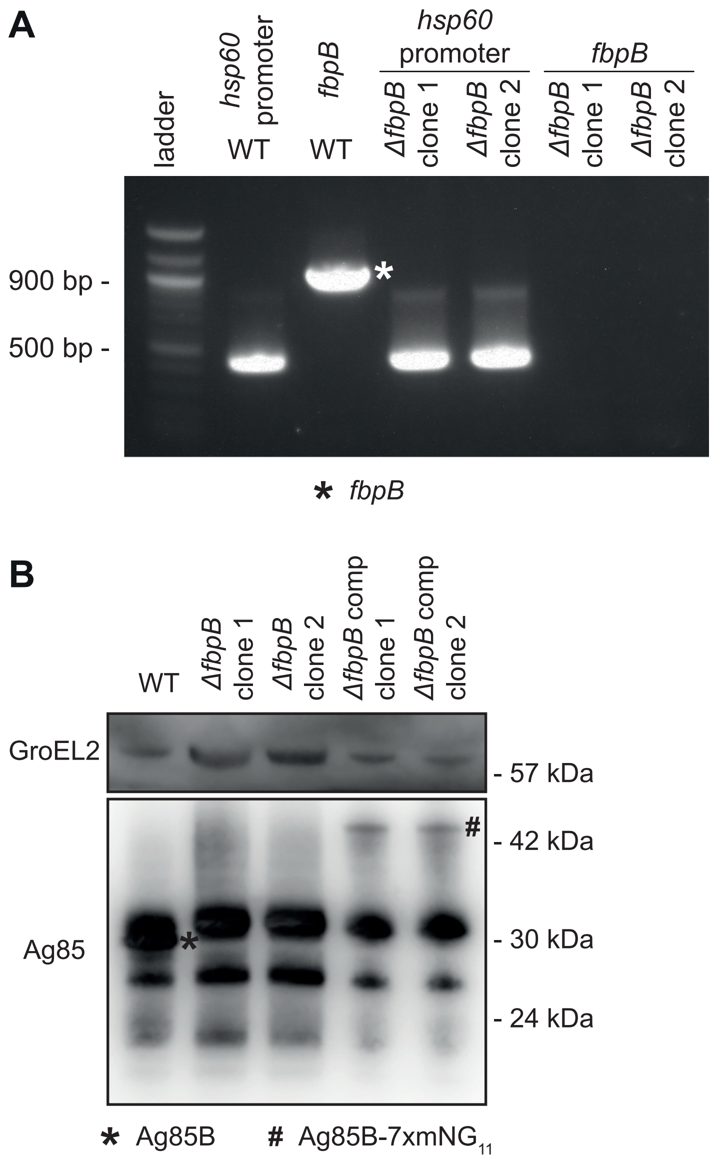

Supp. Figure 7

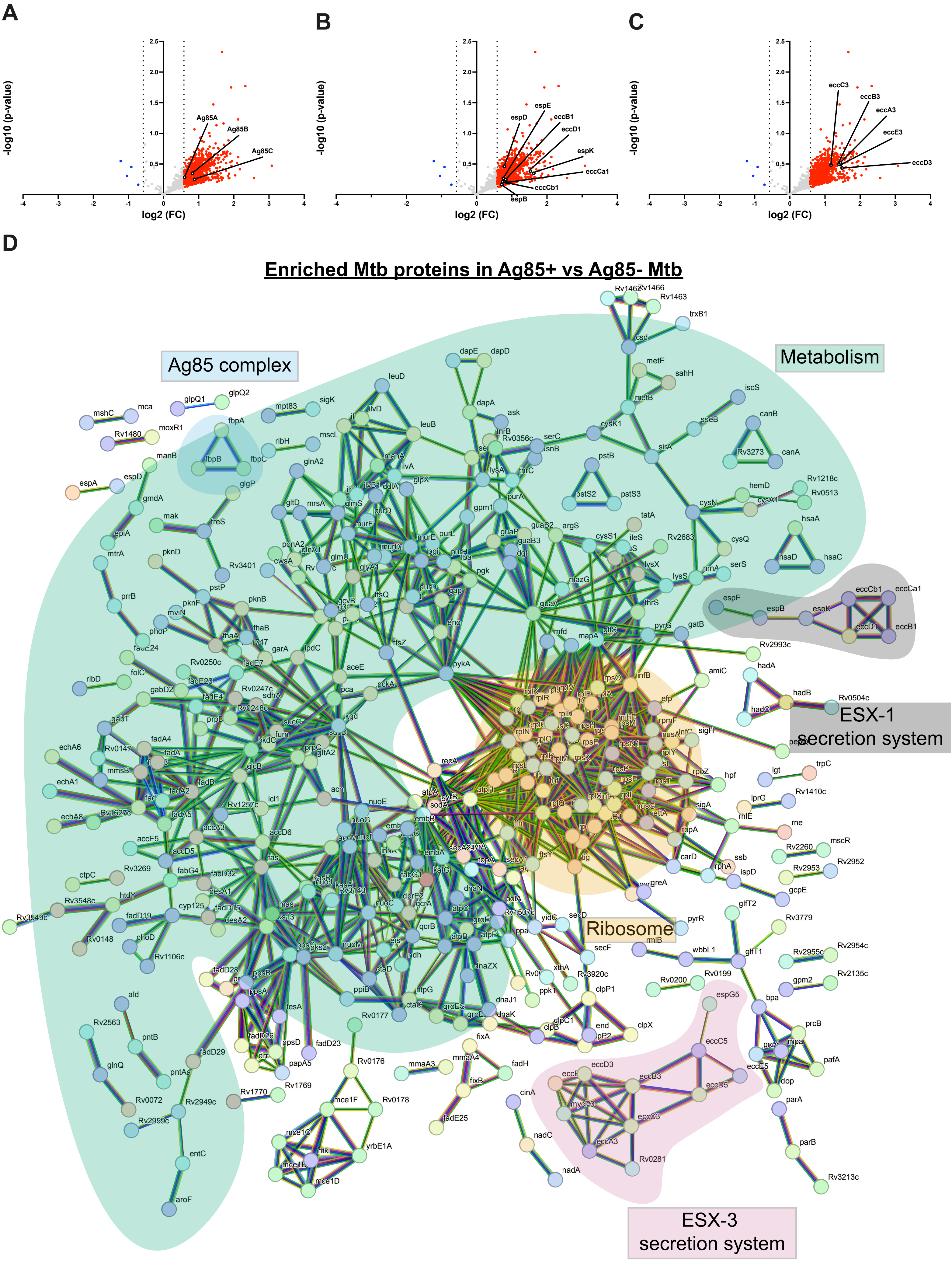

Supp. Figure 8

**A**

### Enriched host protein in Ag85+ vs Ag85- Mtb

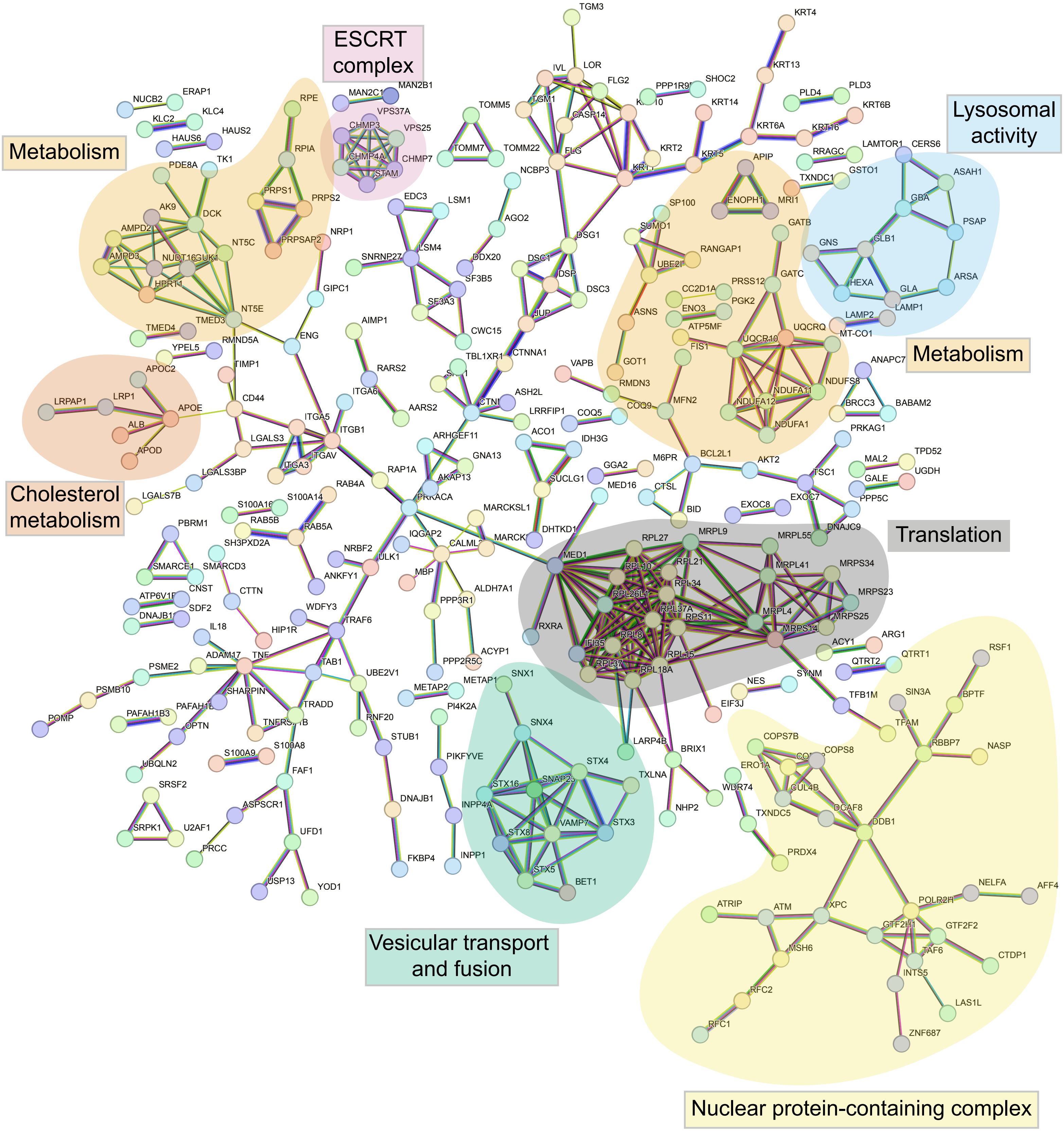

Supp. Figure 9

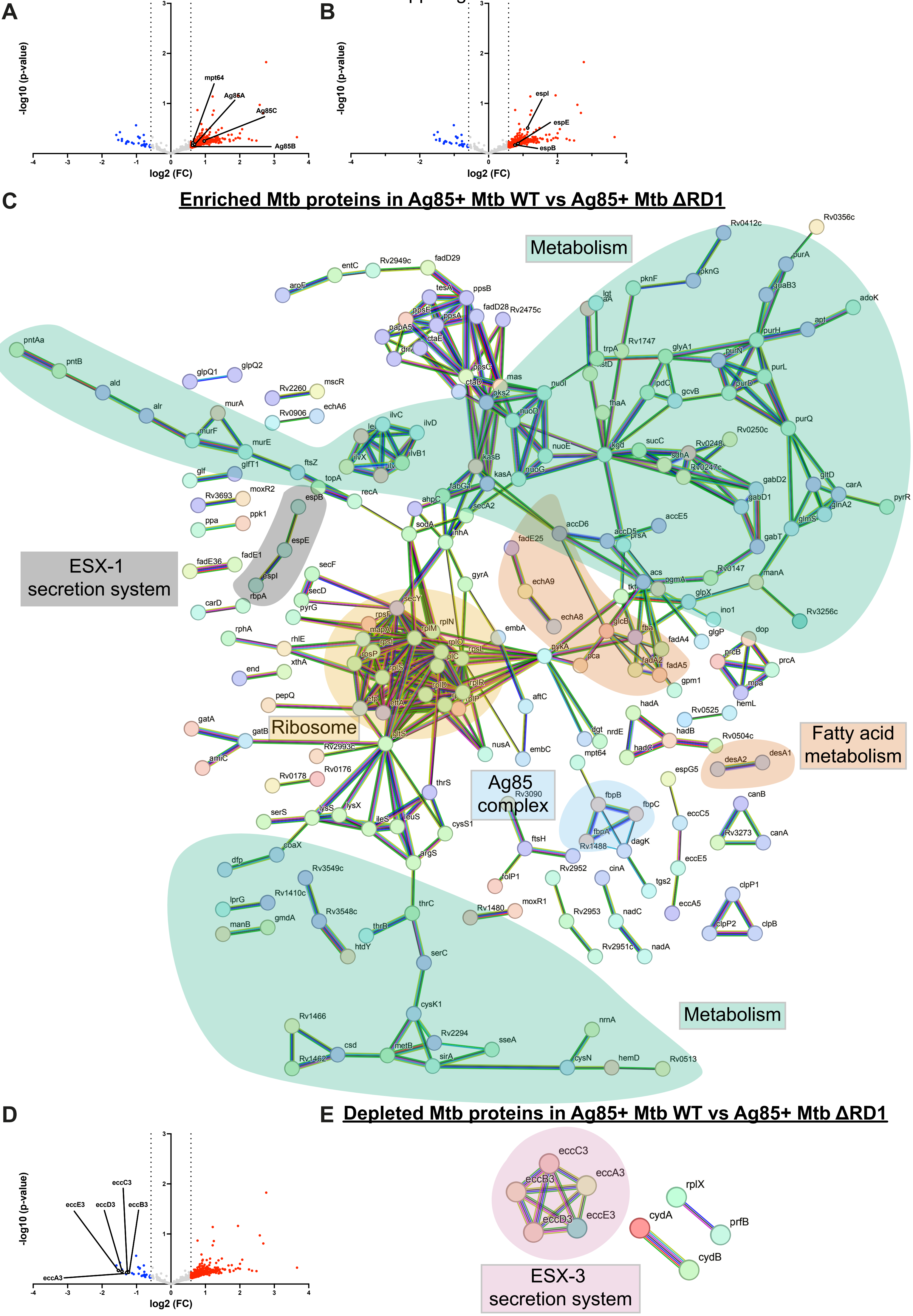

Supp. Figure 10

**A**

### Enriched host proteins in Ag85+ Mtb WT vs Ag85+ Mtb ΔRD1

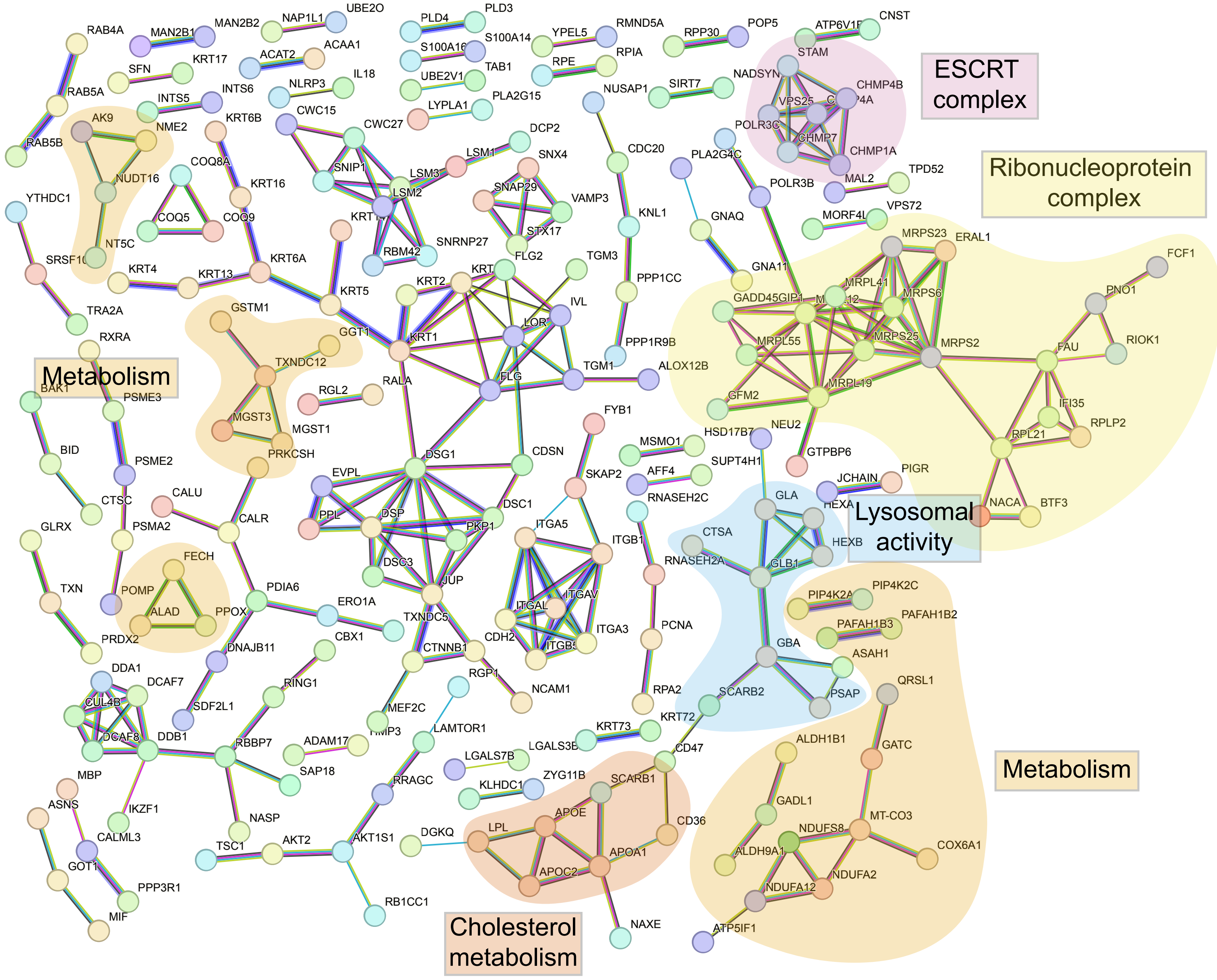
